# Characterizing the rates and patterns of *de novo* germline mutations in coppery titi monkeys (*Plecturocebus cupreus*)

**DOI:** 10.64898/2026.01.15.699688

**Authors:** Cyril J. Versoza, Karen L. Bales, Jeffrey D. Jensen, Susanne P. Pfeifer

## Abstract

Although recent advances in genomics have enabled the high-resolution study of whole genomes, our understanding of one of the key evolutionary processes, mutation, still remains limited. In primates specifically, studies have largely focused on humans and their closest evolutionary relatives, the great apes, as well as a handful of species of biomedical or conservation interest. Yet, as biological variation in mutation rates has been shown to vary across genomic regions, individuals, and species, a greater understanding of the underlying evolutionary dynamics at play will ultimately be illuminated by not only additional sampling across the Order, but also by a greater depth of sampling within-species. To address these needs, we here present the first population-scale genomic resources for a platyrrhine of considerable biomedical interest for both social behavior and neurobiology, the coppery titi monkey (*Plecturocebus cupreus*). Deep whole-genome sequencing of 15 parent-offspring trios, together with a computational *de novo* mutation detection pipeline based on pan-genome graphs, has provided a detailed picture of the sex-averaged mutation rate — 0.63 × 10^-8^ (95% CI: 0.43 × 10^-8^ – 0.90 × 10^-8^) per site per generation — as well as the effects of both sex and parental age on underlying rates, demonstrating a significant paternal age effect. Coppery titi monkey males exhibit long reproductive lifespans, afforded by long-term pair bonding in the species’ monogamous mating system, and our results have demonstrated that individuals reproducing later in life exhibit one of the strongest male mutation biases observed in any non-human primate studied to date. Taken together, this study thus provides an important piece of the puzzle for better comprehending the mutational landscape across primates.

## INTRODUCTION

A platyrrhine native to the north-central neotropical forests of South America, coppery titi monkeys (*Plecturocebus cupreus*; formerly *Callicebus cupreus*; Groves 2005) have emerged as a key primate model for behavioral research. As the species is characterized by long-term socially monogamous mate pairing with an extensive paternal investment in infant care (Mendoza and Mason 1986; Kinzey 1997; Valeggia et al. 1999) — both features uncommon amongst mammals (Lukas and Clutton-Brock 2013) — *P. cupreus* has become a focal point for the investigation of neurobiology, particularly as it pertains to social bonding and behavior important to human health and well-being (see the review of Bales et al. 2017). Notably, and in contrast to other platyrrhines used in biomedical research, the amino acid structure of oxytocin — a hormone that is involved in social bonding and that regulates important aspects of sexual reproduction (e.g., birth and lactation; see the review of Carter 2021) — is conserved between coppery titi monkeys and humans (French et al. 2016), thus facilitating translational studies. For example, relating to clinical studies suggesting that oxytocin could be administered to reduce social impairment in individuals impacted by autism spectrum disorder (see the review of Horta et al. 2020), studies in *P. cupreus* have quantified the effects of this neurohormone on general social behaviors as well as pair bonding (Carter et al. 2020; Bales et al. 2021; Rigney et al. 2022; Arias-del Razo et al. 2022a,b; Zablocki-Thomas et al. 2023a; Witczak et al. 2024). Coppery titi monkeys have similarly been used to study a variety of features related to cognition, associative learning, memory, and the role of vocal communication in social interaction (e.g., Bales et al. 2017; Lau et al. 2020).

Despite this biomedical significance, the evolutionary genomics of *P. cupreus* remains poorly characterized due to a scarcity of genomic resources for the species, greatly limiting the potential for any meaningful genetic studies connecting underlying genotypes with these behavioral phenotypes. As a first step towards mitigating this issue, Pfeifer et al. (2024) recently presented a fully annotated *de novo* genome assembly for the species, combining long-, short-, and linked-read sequencing with Hi-C data to obtain chromosome-length scaffolds. This genomic resource provides a necessary component for the in-depth study of the underlying population-level processes generating, maintaining, and purging variation in this species. The starting point in the characterization of evolutionary processes is mutation, the underlying source of genetic variation. While genetic drift as modulated by population history, natural selection, and recombination are all fundamental for interpreting observed levels and patterns of genetic variation, the rate of new mutation is a key parameter in and of itself, essential for inferring and parameterizing the action of these alternative evolutionary processes as well as for dating the timing of population- or phylogenetic-level events (see the reviews of Pfeifer 2020; Johri et al. 2022). Moreover, and in particular with regards to the great interest in the translational study of behavioral traits in *P. cupreus*, an accurate characterization of the underlying mutational processes will also be crucial for quantifying the role of mutation in health- and disease-related phenotypes (Shendure and Akey 2015).

Germline point mutations are generally thought to originate from uncorrected copying errors during DNA replication, though recent observations have sparked debate (Hahn et al. 2023; Beichman et al. 2024). In primates, sex-specific differences in replication-driven rates are to be expected owing to the larger number of germline cell divisions in males compared to females, leading to the expectation of a greater contribution of paternal relative to maternal *de novo* mutation (DNM; Haldane 1935, 1947; and see Crow 2000). This expectation is widely consistent with observation (Ellegren 2007; Wilson Sayres et al. 2011). Given this, the male mutational burden would be expected to increase with paternal age given the continuation of spermatogenesis throughout adulthood (Ségurel et al. 2014; Goriely 2016). While this paternal age effect has been widely observed, there is also evidence from humans that a male bias already exists at the time of puberty and remains relatively stable thereafter (Jónsson et al. 2017; Gao et al. 2019). Spontaneous, replication-independent DNA damage in gametes owing to extrinsic mutational agents (e.g., ultraviolet radiation and mutagenic chemical agents) thus also likely plays a significant role in these underlying rates (Goldmann et al. 2016; Jónsson et al. 2017; Wu et al. 2020). In addition, biochemical mechanisms of DNA repair efficiency and replication fidelity have been well characterized (see the review of Mohrenweiser et al. 2003), and themselves are significant predictors of genomic rate variation. Perhaps most noteworthy in this regard has been the observation that CpG sites have an order of magnitude higher DNM rate than non-CpG sites in primates studied to date, owing to spontaneous methylation-dependent deamination (Nachman and Crowell 2000; Hwang and Green 2004; Leffler et al. 2013).

Generally speaking, there are two classes of approach for genetically characterizing these mutational processes in long-generation time species that are not amenable to techniques commonly employed in lab-tractable organisms. Indirect approaches involve the counting of neutral divergent sites between closely related species, thereby relying upon Kimura’s (1968, 1983) observation that the rate of neutral divergence is dictated by the rate of neutral mutation. While exceptionally useful, and capable of providing fine-scale mutation rate maps across a genome, these indirect approaches are also accompanied by considerable uncertainty in the underlying assumptions pertaining to, for example, phylogenetic calibration and generation time scaling (see the review of Drake et al. 1998). For this reason, the gold-standard in primate mutation rate inference has remained direct, pedigree-based approaches. Relying upon recent progress in both computational and sequencing technologies, these approaches count observed *de novo* germline mutations occurring in a single generation by comparing the genomes of parents and their offspring (so-called parent-offspring trios; see the review of Pfeifer 2020). While the accurate discrimination between genuine mutations and sequencing errors remains a bioinformatic challenge, several pipelines have been developed for this purpose demonstrating strong performance characteristics (Pfeifer 2021; Bergeron et al. 2022).

Utilizing these techniques, studies over the past decade in particular have greatly increased our knowledge regarding mutation rates in primates, and have highlighted a substantial variation in rates between species (see the reviews of Tran and Pfeifer 2018; Chintalapati and Moorjani 2020). Outside of humans, while much work has been focused upon the great apes for anthropocentric reasons (e.g., Venn et al. 2014; Tatsumoto et al. 2017; Besenbacher et al. 2019; Ghafoor et al. 2023), recent efforts have been made to extend this inference across the primate clade (e.g., to strepsirrhines; Campbell et al. 2021; Versoza et al. 2025; Soni et al. 2025b). Moreover, despite a particular focus in achieving high-quality rate estimation in biomedically relevant species, including baboons (Wu et al. 2020), rhesus macaques (Wang et al. 2020; Bergeron et al. 2021), owl monkeys (Thomas et al. 2018), and marmosets (Yang et al. 2021; Soni et al. 2025a), coppery titi monkeys have yet to be characterized despite being one of the focal research colonies maintained at the U.S. National Primate Research Centers funded by the U.S. National Institutes of Health. In order to address these needs, we here present the first genomic resources for the species at the population scale — a deep whole-genome sequencing of 15 parent-offspring trios — and utilize recent computational pipeline developments to characterize the rates and patterns of *de novo* germline mutations in *P. cupreus*. Given both the relatively large sample size for a non-human primate and the wide range of paternal ages captured (ranging from 3.0 to 18.3 years of age at the time of the offsprings’ birth), this work provides unique insights into both within-species mutation rate variation as well as paternal- and maternal-age effects. These results thus not only provide an important genotypic piece of the puzzle for further understanding this phenotypically well-studied species, but also offer unique family-level resolution of mutational processes as well as an important primate-comparative estimate in this socially-distinctive platyrrhine.

## RESULTS AND DISCUSSION

### Coppery titi monkey pedigrees

We obtained samples from 25 captive coppery titi monkeys (*P. cupreus*) housed at the California National Primate Research Center (CNPRC), at UC Davis, CA. These individuals formed 15 parent-offspring trios within two three-generation and one two-generation pedigrees (Figure 1): (i) one pedigree comprised of a sire and a dam (parental generation, P_0_) that together produced four first-generation (F_1_) offspring (three females and one male), with an additional three second-generation (F_2_) offspring (two females and one male) derived from three of the F_1_ individuals and their respective partners, (ii) one pedigree including a breeding pair who gave birth to three male F_1_ offspring, with an additional F_2_ female sired by one of the F_1_s, and (iii) one pedigree consisting of parents that had four F_1_ offspring (one female and three males). These pedigrees were selected to cover important time points during the aging process of the species. Specifically, while males reach sexual maturity between 15 and 22 months and females between 29 and 32 months of age (Conley et al. 2022), individuals generally do not reproduce until they disperse from their natal family groups between 2.1 and 5.0 years of age (Van Belle et al. 2016). Under captive management, females generally produce their first young at around 3.7 years of age, although substantial variation has been reported, with ages at first reproduction ranging from 2.0 to 6.9 years (Valeggia et al. 1999); however, comparable information from wild individuals remains lacking. In captivity, males and females exhibit a median lifespan of 14.9 and 11.4 years, respectively (Zablocki-Thomas et al. 2023b), though captive individuals frequently survive into their mid-20s (e.g., individuals as old as 26.2 years having been observed at the CNPRC [Zablocki-Thomas et al. 2023b], and an exceptional case of a captive-born individual reaching the age of 35 years was recorded in the species’ studbook [Vermeer and Baumeyer 2022]). Although long-term demographic data records remain sparse, field studies suggest that the species’ maximum lifespan under natural conditions tends to be considerably shorter, typically reaching between 15 and 20 years (de Magalhães and Costa 2009), with survival in the wild constrained by environmental conditions, predation, and disease. In the pedigrees selected for this study, dams gave birth between 3.1 and 18.3 years of age (median age: 6.9 years), with sires ages ranging from 3.0 to 15.6 years (median age: 8.4 years, and see Figure 1 for the parental ages at the time of birth of their offspring), thus encompassing much of the species’ reproductive life span documented in the wild.

**Figure 1.**
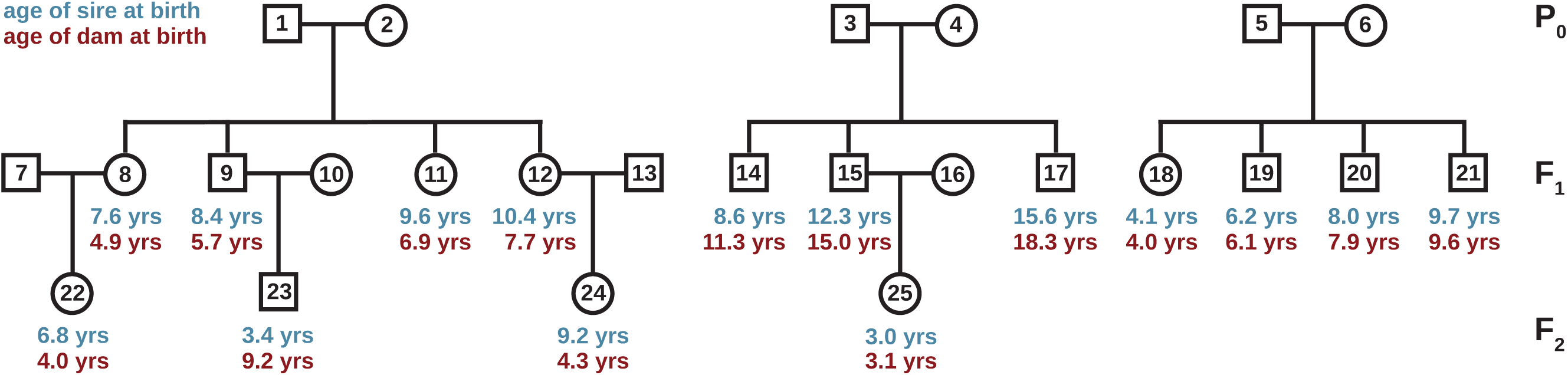
Coppery titi monkey pedigrees. Structure of the two three-generation and one two-generation pedigrees: (**left**) one pedigree comprised of a sire and a dam (parental generation, P_0_) that together produced four first-generation (F_1_) offspring (three females and one male), with an additional three second-generation (F_2_) offspring (two females and one male) derived from three of the F_1_ individuals and their respective partners, (**middle**) one pedigree including a breeding pair who gave birth to three male F_1_ offspring, with an additional F_2_ female sired by one of the F_1_s, and (**right**) one pedigree consisting of parents that had four F_1_ offspring (one female and three males). Male and female individuals are illustrated as squares and circles, respectively. The ages of the sire and dam at the time of birth of their offspring are provided underneath the symbols (shown in blue and red font, respectively).

### Identification of germline DNMs in coppery titi monkeys

We generated whole-genome sequences for the 15 parent–offspring trios, achieving a mean depth of coverage of ∼50× (range: 39.1×–73.4×; Supplementary Table 1). We aligned the quality-controlled reads to the coppery titi monkey genome (GenBank accession number: GCA_040437455.1; Pfeifer et al. 2024) and identified autosomal sites accessible to our study following the Genome Analysis Toolkit (GATK) pipeline for non-model organisms (van der Auwera and O’Connor 2020). As the identification of germline DNMs is sensitive to genotyping errors, we re-genotyped variants discovered with GATK jointly across all individuals using the pan-genome approach implemented in Graphtyper v.2.7.2 (Eggertsson et al. 2017). By reducing the reference bias inherent to linear-reference approaches like GATK, Graphtyper has been shown to lead to increased genotype accuracy, particularly in regions with repetitive or structurally complex loci (Eggertsson et al. 2017). This graph-based pan-genome approach thus allowed us to study DNMs at the genome-wide scale, while avoiding the application of (inherently subjective) sequence-level filtering criteria necessary to eliminate the large number of false positives frequently observed with linear-reference-based approaches (Beal et al. 2012). Although common practice, the reliance on such sequence-level metrics, particularly those that lack a clear analogue for invariant positions, complicates the accurate delineation of the genomic regions that can effectively be interrogated (Pfeifer 2021). As knowledge of this accessible genome is an essential component for estimating per-site mutation rates, differences in filtering strategies can thus lead to considerable variation in mutation rate estimates (Bergeron et al. 2022).

From this re-genotyped call set of 19.2 million autosomal, biallelic single-nucleotide polymorphisms (SNPs; Supplementary Table 2), we identified 995 loci displaying Mendelian inconsistencies across the 15 parent–offspring trios, defined here as sites at which both parents were homozygous for the reference allele and their focal offspring was heterozygous for a non-reference (alternate) allele. To guard against incorrect genotype assignments that could result in false positives, we confirmed the absence of reads supporting the alternate allele in the parents via both the read alignments and the haplotypes locally re-assembled by GATK and Graphtyper. However, guarding against genotyping errors in the offspring is generally more challenging. Although experimental validation of DNMs by PCR amplification and Sanger sequencing is theoretically straightforward, in practice, such approaches are often substantially hampered in non-model organisms for which genomic resources remain scarce or incomplete. For example, fragmented or locally misassembled reference assemblies can complicate primer design, increase the likelihood of non-specific amplification, and lead to elevated assay failure rates, even for genuine variants. These challenges have been well-documented in closely related systems; for instance, a non-human primate study of six parent-offspring trios reported assay failure rates of more than 20% (Venn et al. 2014; and see Bergeron et al. 2022 for discussion). Therefore, we instead implemented a stringent manual curation strategy to evaluate Mendelian-inconsistent sites for genotyping errors following best practices in the field (Bergeron et al. 2022) (for details, see “Identification of germline DNMs”). Following the independent curation of two researchers, 448 of the 995 candidate sites were retained (Supplementary Table 3), with the majority of false positives associated with systematic genotyping errors occurring in the vicinity of homopolymeric tracts (for an example, see Supplementary Figure 1). Multiple independent observations support the interpretation that the DNMs retained after visual inspection represent genuine DNMs rather than technical artefacts: (i) none of the validated DNMs were harbored within genomic regions affected by structural variation (Versoza et al. 2026b) or in close proximity (within 5 bp) of insertions and deletions — genomic contexts that frequently inflate false-positive single-nucleotide calls from short-read data (Sedlazeck et al. 2018), and (ii) tracking the transmission of DNMs across generations, the patterns of inheritance closely matched Mendelian expectations (with average individual transmission rates between 0.41 and 0.57; binomial test *p*-value: 0.5946). These checks thus provide an additional layer of validation as, for example, substantial departures from the expected segregation ratio may indicate undetected technical artefacts and/or the inclusion of early post-zygotic mutations.

### Genomic distribution and mutational signatures of DNMs in coppery titi monkeys

The genomic distribution of DNMs was consistent with chromosomal length (×^2^ = 20.318, df = 21, *p*-value = 0.5012), providing no evidence for chromosome-specific mutation rate heterogeneity in coppery titi monkeys. Mutation rate heterogeneity was, however, observed within individual chromosomes, with 15.0% of DNMs clustering within 1 Mb of another event, suggesting the presence of localized mutational hotspots — an observation in agreement with pedigree-based mutation rate studies of other non-human primates (Campbell et al. 2012; Michaelson et al. 2012; Venn et al. 2014; Francioli et al. 2015). As expected from the composition of the species’ genome, the vast majority of these DNMs occurred in non-coding regions, with intergenic and intronic regions accounting for 76.1% and 17.0% of mutations (Supplementary Figure 2; ×^2^ = 9.6816, df = 6, *p*-value = 0.1387). Out of the 14 DNMs (3.1%) identified within exonic regions, 11 were missense variants of moderate effect (predicted to affect the genes ADHFE1, DDIAS, FMNL2, GVQW1, PDE6C3, and RC3H2) and 3 were synonymous changes of low effect (predicted to affect the genes SYT17 and MTOR). Moreover, 39.3% of DNMs were harbored within annotated repeats, consistent with the overall abundance of repetitive elements in the coppery titi monkey genome (38.7% [Pfeifer et al. 2024]; binominal test *p*-value = 0.8085). Similar to many other eukaryotes, transposable elements represent a large proportion of this repetitive genome (Pfeifer et al. 2024). As transposable elements are highly mutagenic — often disrupting genes, modifying gene expression, and causing genomic rearrangements that negatively impact evolutionary fitness or contribute to genetic disease (Payer and Burns 2019) — many taxa have evolved epigenetic mechanisms to silence their activity (Slotkin and Martienssen 2007). A well-known consequence of such epigenetic modifications is an elevated mutability of methylated CpG dinucleotides that undergo spontaneous methylation-dependent deamination (Hwang and Green 2004; Hodgkinson and Eyre-Walker 2011). Such sites often contribute disproportionately to DNMs; in humans, for example, CpG>TpG mutations give rise to ∼17–19% of all DNMs (Kong et al. 2012; Ségurel et al. 2014). The relative contribution of CpG>TpG mutations in the coppery titi monkey genome (18.0%) falls within this range observed in humans and is similar to that reported in strepsirrhines (17.6%; Versoza et al. 2025; and see the review of Soni et al. 2025c); moreover, the transition-transversion ratio (Ts/Tv) of the identified DNMs (1.75) is statistically similar to that observed in humans (∼2.0 [Kong et al. 2012]; binomial test *p*-value: 0.1761). In contrast, in owl monkeys — the only other platyrrhine for which direct mutation estimates from multiple trios exist to date — the overall contribution of CpG>TpG mutations appears substantially lower (∼12%; Thomas et al. 2018), resulting in significant differences in the mutational spectra between these two species (×^2^ = 25.16, df = 5, *p*-value < 0.0001; Figure 2). Further extending the sequence-context of each DNM by their 5’ and 3’ flanking nucleotides and combining strand complements, we used the observed proportion of the 96 trinucleotide mutational events to study the activity of COSMIC single-base mutational signatures (SBS; Alexandrov et al. 2020). The vast majority of DNMs exhibited SBS5 (74.8%) and SBS1 (11.8%) mutational signatures — an observation consistent with previous studies of the mammalian germline (Rahbari et al. 2016; Spisak et al. 2024). Both SBS5 and SBS1 are thought to accrue in a “clock-like” fashion, with the latter associated with the methylation-mediated deamination of 5-methylcytosine (Alexandrov et al. 2013). In addition, a smaller proportion (∼14%) of SBS6 signatures contribute to the observed DNMs, highlighting the role of defective DNA mismatch repair in the mutational processes governing the evolution of the coppery titi monkey genome.

**Figure 2.**
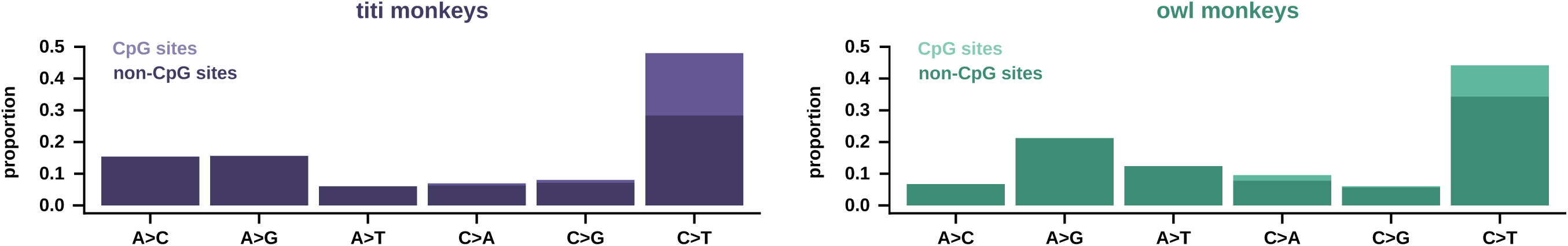
Mutational spectrum of the coppery titi monkey. Mutational spectra of platyrrhines indicating the relative proportion of each mutation type (with reverse complements collapsed): (**left**) mutational spectrum of coppery titi monkey DNMs based on 15 parent-offspring trios (shown in purple; this study) and (**right**), for comparison, owl monkeys — the only other platyrrhine for which direct mutation estimates from multiple trios exist to date (based on 14 parent-offspring trios shown in teal; Thomas et al. 2018).

### Estimation of per-generation germline mutation rates and parental age effects

In order to estimate per-site per-generation germline mutation rates, we first needed to quantify the false negative rate (FNR) of our study. To this end, we followed the simulation-based methodology described in Pfeifer (2017a), in which synthetic DNMs are introduced into the haplotype-resolved reads of the offspring before processing these modified reads with the same computational workflows used to identify DNMs. Based on the fraction of synthetic DNMs missed by our DNM discovery pipeline, we estimated a FNR of 3.18%. Based on the length of the autosomal genome accessible to our study (∼4.8 Gb per trio), and correcting for both false positive and false negative rates, we estimated an average autosomal per-site per-generation point mutation rate of 0.63 × 10^-8^ (95% CI: 0.43 × 10^-8^ – 0.90 × 10^-8^). Inferred mutation rates varied between ∼0.5 × 10^-8^ per base pair per generation (/bp/gen) in individuals born to younger parents (with the earliest birth observed at a parental age of ∼3.0 years) and ∼1.1 × 10^-8^ /bp/gen in individuals born to older parents (with paternal and maternal ages at birth of 15.6 and 18.3 years, respectively) (Figure 3a). These estimates are thus within the range of the average direct per-generation germline mutation rate estimates previously inferred from pedigree-based studies of other primates: 1.05 × 10^-8^ – 1.29 × 10^-8^ /bp in humans based on 100s to 1000s of parent-offspring trios (Francioli et al. 2015; Wong et al. 2016; Jónsson et al. 2017; Maretty et al. 2017), 1.20 × 10^-8^ – 1.26 × 10^-8^ /bp in chimpanzees based on six to seven trios (Venn et al. 2014; Besenbacher et al. 2019), 1.13 × 10^-8^ /bp in gorillas based on two trios (Besenbacher et al. 2019), 1.66 × 10^-8^/bp in orangutan based on a single trio (Besenbacher et al. 2019), 0.58 × 10^-8^ – 0.77 × 10^-8^ /bp in rhesus macaques based on 14–19 trios (Wang et al. 2020; Bergeron et al. 2021), 0.81 × 10^-8^ /bp in owl monkeys based on 14 trios (Thomas et al. 2018), 0.94 × 10^-8^ /bp in green monkeys based on three trios (Pfeifer 2017a), 0.43 × 10^-8^ /bp in common marmosets based on a single trio (Yang et al. 2021), 1.52 × 10^-8^ /bp in gray mouse lemurs based on two trios (Campbell et al. 2021), and 1.1 × 10^-8^ /bp in aye-ayes based on seven trios (Versoza et al. 2025). Given the average parental age of 8.0 years observed in the 15 parent-offspring trios of our study (Supplementary Table 1) — and consistent with the generation time previously reported in the species (Pacifici et al. 2013) — this estimate thus yields an average estimated yearly mutation rate of 0.78 × 10^-9^ /bp. As anticipated from differences in life history traits, the estimated yearly mutation rate for coppery titi monkeys is considerably higher than the rate estimated for humans (∼0.4 × 10^-9^ /bp, assuming an age of puberty ∼13 years and a parental age of conception ∼30 years; Jónsson et al. 2017) but lower than that estimated for owl monkeys (∼1.2 × 10^-9^ /bp, assuming an age of puberty ∼1 year and a parental age of conception ∼6.5 years; Thomas et al. 2018).

**Figure 3.**
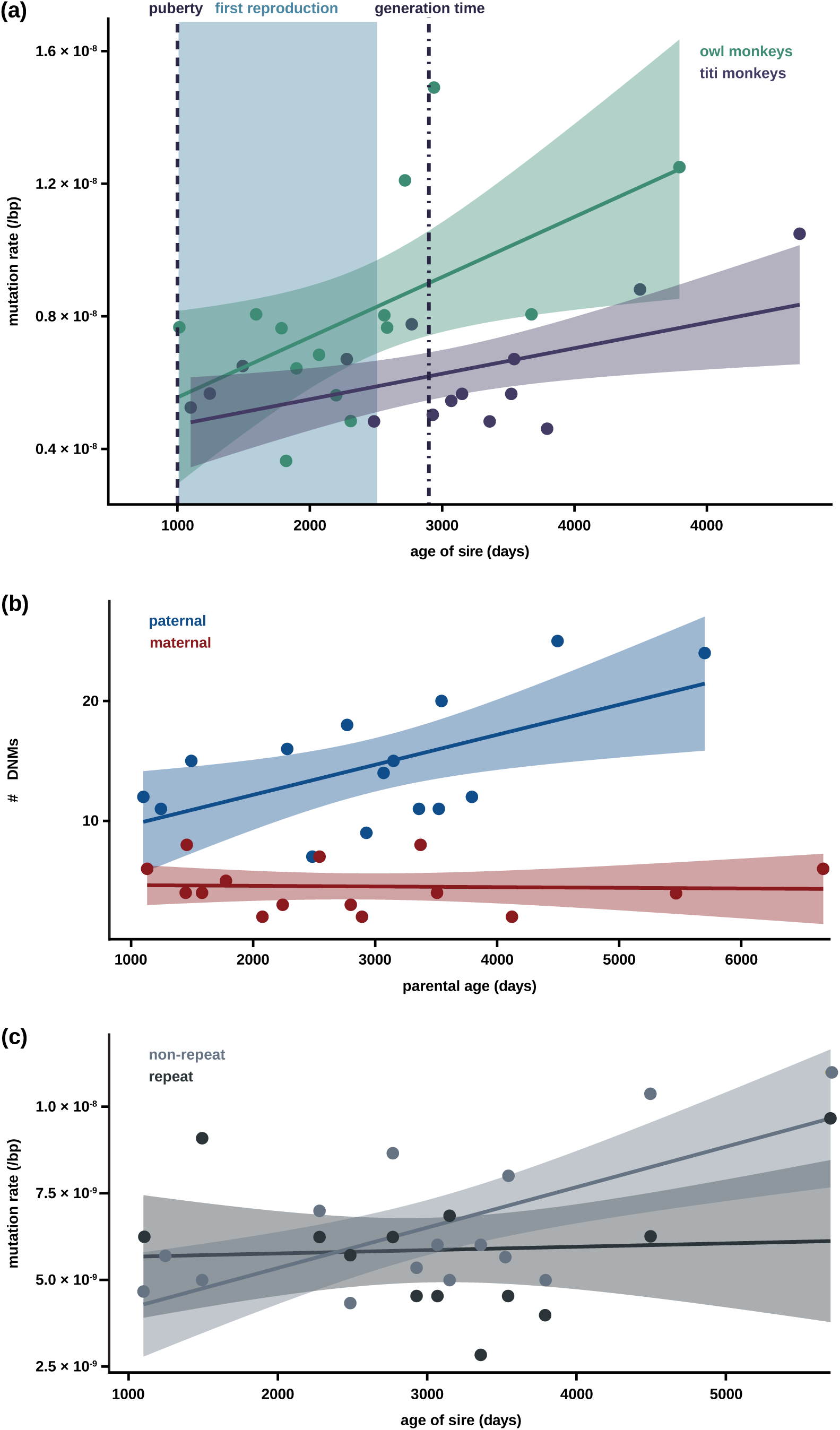
Mutation rate estimate of the coppery titi monkey. Per-site per-generation mutation rate estimates of platyrrhines. (a) Relationship between the paternal age at birth (in days) and the per-site per-generation mutation rate in coppery titi monkeys based on 15 parent-offspring trios (shown in purple; this study) and, for comparison, owl monkeys — the only other platyrrhine for which direct mutation estimates from multiple trios exist (based on 14 parent-offspring trios shown in teal; Thomas et al. 2018). Linear regression and 95% confidence intervals are shown as solid lines and shaded areas, respectively. Dashed and dot-dashed lines indicate the time of sexual maturity and the generation time in coppery titi monkeys, respectively. The age range at first reproduction in coppery titi monkeys is shown as a light blue shaded box. (b) Relationship between parental age at birth (in days) and the number of DNMs for which the parent-of-origin could be determined (with maternal DNMs shown in red and paternal DNMs shown in blue). (c) Relationship between the paternal age at birth (in days) and the per-site per-generation mutation rate in coppery titi monkeys outside and within of repetitive regions (light and dark gray, respectively).

As coppery titi monkeys are characterized by long-term socially monogamous mate pairing, maternal and paternal ages showed a significantly positive correlation (Spearman’s ρ = 0.66, *p*-value: 0.009). In order to study the sex-specific impact of parental ages on mutation rates in the species, we thus first determined the parent-of-origin of the DNMs using read-tracing, which assigned 64.0% of DNMs per trio on average to a parental haplotype (range: 45.8%–75.0%). Based on these DNMs with known parent-of-origin (Supplementary Table 3), we observed a significant paternal age effect on germline mutation rates, with the rate of paternally-derived DNMs increasing by ∼18% per 1,000 days of paternal age (Poisson regression; *p*-value = 0.003); in contrast, no evidence of a maternal age effect was observed in the species (*p*-value = 0.88) (Figure 3b). Notably, the strength of the paternal age effect depends on the genomic background (Figure 3c), and is only statistically significant for non-repetitive genomic regions (*p*-value _non-repeat_ = 0.00439 vs *p*-value _repeat_ = 0.21). These observations are consistent with a male-driven mutational process (though note that maternal age effects tend to be more subtle in primates [Goldmann et al. 2016; Wong et al. 2016; Jónsson et al. 2017] and thus may not be detectable at this sample size).

The average male mutation bias observed in the coppery titi monkey trios is 3.9, consistent with previous estimates in humans (3.1–3.9; Jónsson et al. 2017). Notably however, coppery titi monkeys reproducing later in life show a considerably stronger male bias (up to 7.5) — one of the strongest male mutation biases observed in any non-human primate studied to date (∼4.4, ∼2.0, and ∼4.1 in chimpanzees, gorillas, and orangutans, respectively [Besenbacher et al. 2019], ∼3.0 in rhesus macaques [Wang et al. 2020], ∼3.2 in baboons [Wu et al. 2020], ∼2.1 in owl monkeys [Thomas et al. 2018], ∼2.7 in aye-ayes [Versoza et al. 2025], and ∼1.2 in gray mouse lemurs [Campbell et al. 2021]). This likely reflects a combination of species-specific differences in generation time and life history — in particular the long reproductive lifespan afforded by long-term pair bonding in the species’ monogamous mating system — as well as patterns of germline division. With regards to the latter, no empirical estimates of spermatogonial stem cell division rates yet exist for platyrrhines but differences from the rates observed in humans likely contribute to the differences in male mutation bias between the species. For example, assuming that coppery titi monkey males reach sexual maturity around 15 months of age (Conley et al. 2022), gestation lasts around 132 days (de Magalhães and Costa 2009), and spermatogonial stem cell (SSC) divisions are similar to those previously reported for cercopithecoids (∼33 SSC divisions per year post-puberty; Chowdhury and Steinberger 1976), approximately 462 SSC divisions would be expected to have occurred post-puberty at the time of reproduction for the oldest male included in this study. That is, 67.4% more than in humans (assuming a male age of puberty of ∼13 years in humans [Heller and Clermont 1963], ∼23 SSC divisions per year post-puberty [Drost and Lee 1995], and an average age of reproduction of ∼25 years in humans [Fenner 2005]). As smaller species tend to have higher rates of SSC divisions, the actual difference is presumably even greater and may thus potentially account for the considerably stronger male bias observed in older coppery titi monkeys.

## CONCLUSION

To date, insights into the rates and patterns of *de novo* germline mutation governing the evolution of primate genomes remain limited to a few species of anthropocentric or biomedical interest. Moreover, even amongst comparatively well-studied non-human primates, estimation is frequently based on a handful of trios, thus preventing any insight into, for example, family-level mutation rate variation. Studying 15 parent-offspring trios sampled across the long reproductive lifespan of coppery titi monkeys, we here provide the first direct mutation rate estimates for this platyrrhine of considerable biomedical interest for both social behavior and neurobiology. While the species’ sex-averaged mutation rate falls within the range of that of other primates, substantial variation exists depending on parental age, by and large driven by a strong paternal age effect and male mutation bias. The mutational signatures observed in the coppery titi monkey genome suggest that, similar to humans, most mutations accrue in a “clock-like” manner over time, further highlighting the importance of encompassing a species’ reproductive life span when studying mutation rates within (and between) species and incorporating this sex- and age-specific variation into evolutionary models for dating the timing of population- or species-level events.

## MATERIALS AND METHODS

### Animal subjects

Animals were maintained at the CNPRC. This study was performed in compliance with all regulations regarding the care and use of captive primates, including the NIH Guidelines for the Care and Use of Animals and the American Society of Primatologists’ Guidelines for the Ethical Treatment of Nonhuman Primates. Procedures were approved by the UC-Davis Institutional Animal Care and Use Committee (protocol 22523).

### Whole-genome sequencing

We collected blood samples from 25 captive coppery titi monkeys (*Plecturocebus cupreus*; 13 males and 12 females) spanning two three-generation and one two-generation pedigrees (Figure 1). We isolated high-molecular weight genomic DNA from the samples using either the PAXgene Blood DNA System or the QIAamp DNA Mini Kit (Qiagen, Hilden, Germany). We quantified DNA yields with an Invitrogen Qubit Fluorometer (Thermo Fisher Scientific, Waltham, MA, USA) and evaluated DNA quality by agarose gel electrophoresis. For each individual, we constructed 150 bp paired-end sequencing libraries, following the Illumina TruSeq DNA PCR-Free protocol (Illumina, San Diego, CA, USA). We quantified the libraries using Qubit fluorometry and real-time PCR, and evaluated them for fragment size distribution using a Bioanalyzer (Agilent Technologies, Santa Clara, CA, USA), before generating high-coverage, whole-genome sequencing data on an Illumina NovaSeq 6000 platform (Supplementary Table 1).

### Read pre-processing

To remove experimental artefacts and ensure accurate read alignment and variant calling (Pfeifer 2017b), we processed the sequencing data with fastp v.0.24.0 (Chen et al. 2018), enabling the automatic detection and removal of adapter sequences from the paired-end reads (*--detect_adapter_for_pe*). By default, fastp also detects and trims polyG tails from Illumina NovaSeq reads; moreover, the software applies a built-in filtering procedure, discarding any reads in which more than 40% of bases exhibit Phred quality scores below Q15, that contain more than five undetermined nucleotides (Ns), or that are shorter than 15 bp after trimming.

### Read alignment

We aligned the filtered reads to the NCBI reference genome assembly for the species, PleCup_hybrid (GenBank accession number: GCA_040437455.1; Pfeifer et al. 2024), using *fq2bam*, the GPU-accelerated version of BWA-MEM (Li 2013), deployed within the NVIDIA Parabricks v.4.4.0-1 software suite (Zhu et al. 2025), specifying the *-M* flag to mark shorter split alignments as secondary. We then combined aligned reads originating from different sequencing runs of the same individual with the *MergeSamFiles* function implemented in GATK v.4.2.6.1 (van der Auwera and O’Connor 2020) and marked duplicates with Parabricks’ *markdup*.

### Alignment post-processing

As no experimentally validated set of polymorphic sites yet exists for coppery titi monkeys, we followed the developer-recommended bootstrapping procedure to generate our own high-confidence variant set to iteratively train GATK’s base quality score recalibration (BQSR) model. Briefly, we performed an initial round of variant calling without BQSR in gVCF mode (*--gvcf*) per individual using Parabricks’ *haplotypecaller* (i.e., the GPU-accelerated version of GATK’s *HaplotypeCaller*) on the duplicate-marked reads, requiring a minimum mapping quality of 40 (*--minimum-mapping-quality* 40) and disabling PCR indel error modeling (*-pcr-indel-model* NONE). We then combined gVCFs across all individuals using GATK *CombineGVCFs* and performed joint genotyping to generate a preliminary, multi-individual variant call set using *GenotypeGVCFs*. To derive a provisional set of high-confidence variants suitable for bootstrapping, we applied hard filtering to autosomal, biallelic SNPs genotyped in all individuals based on GATK-recommended annotations and empirically determined thresholds that preserved appropriate transition–transversion ratios following previous studies (e.g., Auton et al. 2012). Specifically, using BCFtools *filter* v.1.14 (Danecek et al. 2021), we excluded SNPs with a quality-by-depth (QD) ratio below 10, a Fisher Strand (FS) test value larger than 5, a Symmetric Odds Ratio (SOR) test value larger than 1.5, a rank sum test value for mapping qualities of reads supporting the reference vs the alternate allele (MQRankSum) below −12.5, a rank sum test value for the relative positioning of the reference vs the alternate allele within reads (ReadPosRankSum) below −8.0, a genotype quality (GQ) below 60, or a depth (DP) of less than half or greater than twice of an individual’s autosomal average coverage. This filtered call set was then treated as a temporary “known sites” resource for recalibration. Using this bootstrapped variant set, we performed BQSR (Parabricks’ *bqsr*) to model systematic biases in base quality scores associated with machine cycles and sequence context, and applied the recalibration to the duplicate-marked alignments (*applybqsr*). We assessed convergence of the bootstrapping procedure by confirming stability of recalibration model parameters and variant quality metrics between successive iterations.

### Variant calling and genotyping

For each individual, we called variant and invariant autosomal sites on the BQS-recalibrated reads using the GATK *HaplotypeCaller* in base pair resolution mode (*-ERC* BP_RESOLUTION), requiring a minimum mapping quality of 40 (*--minimum-mapping-quality* 40) and disabling PCR indel error modeling (*--pcr-indel-model* NONE). We then combined the resulting gVCFs across all individuals using *CombineGVCFs* and jointly genotyped all sites (*-all-sites*) to generate a multi-individual call set using *GenotypeGVCFs*. To improve genotyping accuracy, we re-genotyped biallelic SNPs discovered with GATK using Graphtyper v.2.7.2 (Eggertsson et al. 2017) and limited our final dataset to high-confidence sites that passed all built-in filters and exhibited genotype information for all individuals (Supplementary Table 2).

### Identification of germline DNMs

Using BCFtools *view* v.1.14 (Danecek et al. 2021), we identified Mendelian-inconsistent sites in the genomes of the 15 parent-offspring trios by selecting loci at which both parents were homozygous for the reference allele while their offspring was heterozygous for the alternate (non-reference) allele; additionally, we required that none of the unrelated individuals in the dataset carried the alternate allele. To guard against incorrect genotype assignments of the parents, we confirmed the absence of reads supporting the alternate allele in both the read alignments (using BCFtool *mpileup*) and the haplotypes locally re-assembled by GATK and Graphtyper. Following best practices in the field (Bergeron et al. 2022) to exclude false positives and validate genuine DNMs, two researchers independently evaluated the read-support of the parental and filial genotypes at each Mendelian-inconsistent site using Integrated Genomics Viewer (IGV) v.2.16.1 (Thorvaldsdóttir et al. 2012) visualizations, discarding any candidates that showed evidence of technical artefacts (see Figure 4 in Pfeifer 2017b for illustrative examples).

To assess the quality of the final dataset (Supplementary Table 3), we then evaluated the validated DNMs for their proximity to genomic regions affected by structural variation (using the structural variant catalogue of Versoza et al. 2026b) as well as insertions and deletions (using the indels identified in this study) given that these regions can pose challenges for short-read alignment (Sedlazeck et al. 2018), which in turn can give rise to false-positive single-nucleotide calls (Pfeifer 2017b). Moreover, to assess biological plausibility, we examined the transmission patterns of the validated DNMs observed in the four F_1_ individuals with F_2_ progeny. Under Mendel’s Laws of Inheritance, a genuine heterozygous DNM is expected to be passed on to an offspring with a probability of 0.5 (Mendel 1866); we tested whether the observed average transmission rates deviated from this expectation by applying a Fisher’s exact test implemented in R v.4.2.2 (R Core Team 2022).

### Parent-of-origin assignment of DNMs

Following earlier work in other primates (Goldmann et al. 2016; Jónsson et al. 2017), we assigned the parent-of-origin of the DNMs detected in the genomes of the 15 parent-offspring trios using read-tracing. To this end, we searched the 1kb-regions surrounding each DNM for phase-informative (heterozygous) sites located either on the same (or paired) read or linked to the same haplotype than the DNM using the approaches implemented in the Parent Of Origin Haplotype Annotator (POOHA: https://github.com/besenbacher/POOHA; Maretty et al. 2017; Besenbacher et al. 2019) and Unfazed v.1.0.2 (Belyeu et al. 2021).

### Estimation of the per-generation mutation rate

We estimated the autosomal per-site per-generation point mutation rate *μ* as *μ* = # *DNMs*⁄(2 × *CG* × (1 − *FNR*)), where # *DNMs* is the number of validated DNMs, *CG* is the autosomal genome accessible to our study, and *FNR* is the false negative rate of our study. We calculated 95% confidence intervals assuming a Poisson distribution.

To quantify the FNR of our study, we followed the simulation-based methodology described in Pfeifer (2017a), in which synthetic DNMs are introduced into the haplotype-resolved reads of the offspring. To this end, we first reconstructed the haplotypes present in each trio using the pedigree-aware phaser (*--ped*) implemented in WhatsHap *phase* v.2.3 (Patterson et al. 2015; Garg et al. 2016) which integrates read-tracing with genetic phasing, assuming a genome-wide recombination rate of 1.02 cM/Mb, as previously estimated for the species (Versoza et al. 2026a). We then introduced 1,000 synthetic DNMs at randomly selected genomic positions in the phased reads of the offspring using BAMSurgeon *addsnp.py* v.1.4.1 (Ewing et al. 2015). To ensure that synthetic DNMs closely resembled genuine heterozygous sites, we restricted the maximum minor allele frequency of nearby linked polymorphisms to 0.1 (*-s* 0.1). Under these conditions, BAMSurgeon successfully inserted 566 synthetic DNMs. We validated that the patterns of allele balance of these synthetic DNMs closely matched those observed at heterozygous sites (based on loci where each parent was homozygous for a different allele and their offspring was heterozygous; Supplementary Figure 3) before processing the modified reads using the same workflows to identify DNMs described above, recovering 548 of the synthetic 566 DNMs. Based on the fraction of synthetic DNMs not recovered in this call set, we estimated an overall FNR of 3.18% for our DNM discovery pipeline.

### Characterization of the genomic distribution and mutational signatures of DNMs

We classified the validated DNMs by their genomic context based on the gene models available for the coppery titi monkey genome (GenBank accession number: GCA_040437455.1; Pfeifer et al. 2024) using ANNOVAR (release 2020-06-08; Wang et al. 2010) and predicted their functional impact using SnpEff v.5.2 (Cingolani et al. 2012). In order to establish an appropriate null expectation for genomic localization, we applied the same annotation pipeline to the full set of autosomal sites that were genotyped across all individuals (Supplementary Table 2). We then performed a chi-squared (×^2^) goodness-of-fit test to evaluate DNM enrichment in each category relative to the genome-wide composition.

We also classified the validated DNMs according to their specific mutational type, assigned with respect to the coppery titi monkey genome (GenBank accession: GCA_040437455.1; Pfeifer et al. 2024), distinguishing A>C, A>T, C>A, and C>G transversions as well as A>G and C>T transitions (with the latter further subdivided into CpG-contexts and non-CpG contexts), and used the relative frequencies of these classes to characterize the species’ mutational spectrum. Using a ×^2^ goodness-of-fit test, we compared the mutational spectrum of coppery titi monkeys to that of the only other platyrrhine for which direct mutation estimates from multiple trios exist to date, the owl monkey (Thomas et al. 2018). We then extended the sequence-context of each DNM by including information regarding their 5’ and 3’ flanking nucleotides and combined strand complements in order to generate a matrix of 96 trinucleotide mutational events. We re-scaled this matrix by the number of trinucleotide mutational opportunities in the coppery titi monkey genome and adjusted the ratios to those observed in humans (GRCh38 genome build) to account for lineage-specific nucleotide composition. Based on these frequencies, we inferred mutational signature activity using SigProfilerAssignment *cosmic_fit* v.1.1.1 (Díaz-Gay et al. 2023).

## ACKNOWLEDGEMENTS

We would like to thank the California National Primate Research Center for providing the coppery titi monkey samples used in this study. DNA extraction, library preparation, and Illumina sequencing were conducted at the DNA Technologies and Expression Analysis Core at the UC Davis Genome Center (supported by NIH Shared Instrumentation Grant 1S10OD010786-01) and Novogene (Sacramento, CA, USA). Computations were performed on the Sol supercomputer at Arizona State University (Jennewein et al. 2023).

## REFERENCES

Alexandrov LB, Kim J, Haradhvala NJ, Huang MN, Tian Ng AW, Wu Y, Boot A, Covington KR, Gordenin DA, Bergstrom EN, et al. 2020. The repertoire of mutational signatures in human cancer. Nature. 578(7793):94–101.

Alexandrov LB, Nik-Zainal S, Wedge DC, Aparicio SA, Behjati S, Biankin AV, Bignell GR, Bolli N, Borg A, Børresen-Dale AL, et al. 2013. Signatures of mutational processes in human cancer. Nature. 500(7463):415–421.

Arias-del Razo R, Velasco Vazquez ML, Turcanu P, Legrand M, Floch M, Weinstein TAR, Goetze LR, Freeman SM, Baxter A, Witczak LR, et al. 2022a. Long term effects of chronic intranasal oxytocin on adult pair bonding behavior and brain glucose uptake in titi monkeys (*Plecturocebus cupreus*). Horm Behav. 140:105126.

Arias-del Razo R, Velasco Vazquez ML, Turcanu P, Legrand M, Lau AR, Weinstein TAR, Goetze LR, Bales KL. 2022b. Effects of chronic and acute intranasal oxytocin treatments on temporary social separation in adult titi monkeys (*Plecturocebus cupreus*). Front Behav Neurosci. 16:877631.

Auton A, Fledel-Alon A, Pfeifer S, Venn O, Ségurel L, Street T, Leffler EM, Bowden R, Aneas I, Broxholme J, et al. 2012. A fine-scale chimpanzee genetic map from population sequencing. Science. 336(6078):193–198.

Bales KL, Ardekani CS, Baxter A, Karaskiewicz CL, Kuske JX, Lau AR, Savidge LE, Sayler KR, Witczak LR. 2021. What is a pair bond? Horm. Behav. 136:105062.

Bales KL, Arias-del Razo R, Conklin QA, Hartman S, Mayer HS, Rogers FD, Simmons TC, Smith LK, Williams A, Williams DR, et al. 2017. Titi monkeys as a novel non-human primate model for the neurobiology of pair bonding. Yale J Biol Med. 90(3):373–387.

Beal MA, Glenn TC, Somers CM. 2012. Whole genome sequencing for quantifying germline mutation frequency in humans and model species: cautious optimism. Mutat Res. 750(2):96–106.

Beichman AC, Zhu L, Harris K. 2024. The evolutionary interplay of somatic and germline mutation rates. Annu Rev Biomed Data Sci. 7(1):83–105.

Belyeu JR, Sasani TA, Pedersen BS, Quinlan AR. 2021. Unfazed: parent-of-origin detection for large and small *de novo* variants. Bioinformatics. 37(24):4860–4861.

Bergeron LA, Besenbacher S, Bakker J, Zheng J, Li P, Pacheco G, Sinding MS, Kamilari M, Gilbert MTP, Schierup MH, et al. 2021. The germline mutational process in rhesus macaque and its implications for phylogenetic dating. GigaScience. 10(5):giab029.

Bergeron LA, Besenbacher S, Turner T, Versoza CJ, Wang RJ, Price AL, Armstrong E, Riera M, Carlson J, Chen H-Y, et al. 2022. The Mutationathon highlights the importance of reaching standardization in estimates of pedigree-based germline mutation rates. Elife. 11:e73577.

Bergeron LA, Besenbacher S, Zheng J, Li P, Bertelsen MF, Quintard B, Hoffman JI, Li Z, St Leger J, Shao C, et al. 2023. Evolution of the germline mutation rate across vertebrates. Nature. 615(7951):285–291.

Besenbacher S, Hvilsom C, Marques-Bonet T, Mailund T, Schierup MH. 2019. Direct estimation of mutations in great apes reconciles phylogenetic dating. Nat Ecol Evol. 3(2):286–292.

Campbell CD, Chong JX, Malig M, Ko A, Dumont BL, Han L, Vives L, O’Roak BJ, Sudmant PH, Shendure J, et al. 2012. Estimating the human mutation rate using autozygosity in a founder population. Nat Genet. 44(11):1277–1281.

Campbell CR, Tiley GP, Poelstra JW, Hunnicutt KE, Larsen PA, Lee HJ, Thorne JL, Dos Reis M, Yoder AD. 2021. Pedigree-based and phylogenetic methods support surprising patterns of mutation rate and spectrum in the gray mouse lemur. Heredity (Edinb). 127(2):233–244.

Carter CS. 2021. Oxytocin and love: myths, metaphors and mysteries. Compr Psychoneuroendocrinol. 9:100107.

Carter CS, Kenkel WM, MacLean EL, Wilson SR, Perkeybile AM, Yee JR, Ferris CF, Nazarloo HP, Porges SW, Davis JM, et al. 2020. Is oxytocin “nature’s medicine”? Pharmacol Rev. 72(4):829–861.

Chen S, Zhou Y, Chen Y, Gu J. 2018. fastp: an ultra-fast all-in-one FASTQ preprocessor. Bioinformatics. 34(17):i884–i890.

Chintalapati M, Moorjani P. 2020. Evolution of the mutation rate across primates. Curr Opin Genet Dev. 62:58–64.

Chowdhury AK, Steinberger E. 1976. A study of germ cell morphology and duration of spermatogenic cycle in the baboon, *Papio anubis*. Anat Rec. 185(2):155–169.

Cingolani P, Platts A, Wang LL, Coon M, Nguyen T, Wang L, Land SJ, Lu X, Ruden DM. 2012. A program for annotating and predicting the effects of single nucleotide polymorphisms, SnpEff: SNPs in the genome of *Drosophila melanogaster* strain w1118; iso-2; iso-3. Fly (Austin). 6(2):80–92.

Conley AJ, Berger T, Del Razo RA, Cotterman RF, Sahagún E, Goetze LR, Jacob S, Weinstein TAR, Dufek ME, Mendoza SP, et al. 2022. The onset of puberty in colony-housed male and female titi monkeys (*Plecturocebus cupreus*): possible effects of oxytocin treatment during peri-adolescent development. Horm Behav. 142:105157.

Crow JF. 2000. The origins, patterns and implications of human spontaneous mutation. Nat Rev Genet. 1(1):40–47.

Danecek P, Bonfield JK, Liddle J, Marshall J, Ohan V, Pollard MO, Whitwham A, Keane T, McCarthy SA, Davies RM. 2021. Twelve years of SAMtools and BCFtools. GigaScience. 10(2):giab008.

de Magalhães JP, Costa J. 2009. A database of vertebrate longevity records and their relation to other life-history traits. J Evol Biol. 22(8):1770–1774.

Díaz-Gay M, Vangara R, Barnes M, Wang X, Islam SMA, Vermes I, Duke S, Narasimman NB, Yang T, Jiang Z, et al. 2023. Assigning mutational signatures to individual samples and individual somatic mutations with SigProfilerAssignment. Bioinformatics. 39(12):btad756.

Drake JW, Charlesworth B, Charlesworth D, Crow JF. 1998. Rates of spontaneous mutation. Genetics. 148(4):1667–1686.

Drost JB, Lee WR. 1995. Biological basis of germline mutation: comparisons of spontaneous germline mutation rates among Drosophila, mouse, and human. Environ Mol Mutagen. 25(Suppl 26):48–64.

Eggertsson HP, Jonsson H, Kristmundsdottir S, Hjartarson E, Kehr B, Masson G, Zink F, Hjorleifsson KE, Jonasdottir A, Jonasdottir A, et al. 2017. Graphtyper enables population-scale genotyping using pangenome graphs. Nat Genet. 49(11):1654–1660.

Ellegren H. 2007. Characteristics, causes and evolutionary consequences of male-biased mutation. Proc Biol Sci. 274(1606):1–10.

Ewing AD, Houlahan KE, Hu Y, Ellrott K, Caloian C, Yamaguchi TN, Bare JC, P’ng C, Waggott D, Sabelnykova VY, et al. 2015. Combining tumor genome simulation with crowdsourcing to benchmark somatic single-nucleotide-variant detection. Nat Methods. 12(7):623–630.

Fenner JN. 2005. Cross-cultural estimation of the human generation interval for use in genetics-based population divergence studies. Am J Phys Anthropol. 128(2):415–423.

Francioli LC, Polak PP, Koren A, Menelaou A, Chun S, Renkens I; Genome of the Netherlands Consortium; van Duijn CM, Swertz M, Wijmenga C, et al. 2015. Genome-wide patterns and properties of *de novo* mutations in humans. Nat Genet. 47(7):822–826.

French JA, Taylor JH, Mustoe AC, Cavanaugh J. 2016. Neuropeptide diversity and the regulation of social behavior in New World primates. Front Neuroendocrinol. 42:18–39.

Gao Z, Moorjani P, Sasani TA, Pedersen BS, Quinlan AR, Jorde LB, Amster G, Przeworski M. 2019. Overlooked roles of DNA damage and maternal age in generating human germline mutations. Proc Natl Acad Sci USA. 116(19):9491–9500.

Garg S, Martin M, Marschall T. 2016. Read-based phasing of related individuals. Bioinformatics. 32(12):i234–i242.

Ghafoor S, Santos J, Versoza C, Jensen JD, Pfeifer SP. 2023. The impact of sample size and population history on observed mutational spectra: a case study in human and chimpanzee populations. Genome Biol Evol. 15(3):evad019.

Goldmann JM, Wong WS, Pinelli M, Farrah T, Bodian D, Stittrich AB, Glusman G, Vissers LE, Hoischen A, Roach JC, et al. 2016. Parent-of-origin-specific signatures of *de novo* mutations. Nat Genet. 48(8):935–939.

Goriely A. 2016. Decoding germline *de novo* point mutations. Nat Genet. 48(8):823–824.

Groves CP. 2005. Species of *Callicebus* (*Callicebus*) *cupreus*. In: Wilson DE, Reeder DM, editors. Mammal species of the world: a taxonomic and geographic reference. 3rd ed. Baltimore: John Hopkins University Press. p. 142–143.

Hahn MW, Peña-Garcia Y, Wang RJ. 2023. The ‘faulty male’ hypothesis for sex-biased mutation and disease. Curr Biol. 33(22):R1166–R1172.

Haldane JBS. 1935. The rate of spontaneous mutation of a human gene. J Genet. 83(3):235–244.

Haldane JBS. 1947. The mutation rate of the gene for haemophilia, and its segregation ratios in males and females. Ann Eugen. 13(1):262–271.

Heller CG, Clermont Y. 1963. Spermatogenesis in man: an estimate of its duration. Science. 140(3563):184–186.

Hodgkinson A, Eyre-Walker A. 2011. Variation in the mutation rate across mammalian genomes. Nat Rev Genet. 12(11):756–766.

Horta M, Kaylor K, Feifel D, Ebner NC. 2020. Chronic oxytocin administration as a tool for investigation and treatment: a cross-disciplinary systematic review. Neurosci Biobehav Rev. 108:1–23.

Hwang DG, Green P. 2004. Bayesian Markov chain Monte Carlo sequence analysis reveals varying neutral substitution patterns in mammalian evolution. Proc Natl Acad Sci USA. 101(39):13994–14001.

Jennewein DM, Lee J, Kurtz C, Dizon W, Shaeffer I, Chapman A, Chiquete A, Burks J, Carlson A, Mason N, et al. 2023. The Sol Supercomputer at Arizona State University. In Practice and Experience in Advanced Research Computing 2023: Computing for the Common Good (PEARC’23). Association for Computing Machinery, New York, NY, USA, 296–301.

Johri P, Aquadro CF, Beaumont M, Charlesworth B, Excoffier L, Eyre-Walker A, Keightley PD, Lynch M, McVean G, Payseur BA, et al. 2022. Recommendations for improving statistical inference in population genomics. PLoS Biol. 20(5):e3001669.

Jónsson H, Sulem P, Kehr B, Kristmundsdottir S, Zink F, Hjartarson E, Hardarson MT, Hjorleifsson KE, Eggertsson HP, Gudjonsson SA, et al. 2017. Parental influence on human germline *de novo* mutations in 1,548 trios from Iceland. Nature. 549(7673):519–522.

Kimura M. 1968. Evolutionary rate at the molecular level. Nature. 217(5129):624–626.

Kimura M. 1983. The Neutral Theory of Molecular Evolution. Cambridge (MA): Cambridge University Press.

Kinzey W. 1997. New World primates: ecology, evolution, and behavior. New York: Aldine de Gruyter.

Kong A, Frigge ML, Masson G, Besenbacher S, Sulem P, Magnusson G, Gudjonsson SA, Sigurdsson A, Jonasdottir A, Jonasdottir A, et al. 2012. Rate of *de novo* mutations and the importance of father’s age to disease risk. Nature. 488(7412):471–475.

Lau AR, Clink DJ, Bales KL. 2020. Individuality in the vocalizations of infant and adult coppery titi monkeys (*Plecturocebus cupreus*). Am J Primatol. 82(6):e23134.

Leffler EM, Gao Z, Pfeifer S, Ségurel L, Auton A, Venn O, Bowden R, Bontrop R, Wall JD, Sella G, et al. 2013. Multiple instances of ancient balancing selection shared between humans and chimpanzees. Science. 339(6127):1578–1582.

Li H. 2013.Aligning sequence reads, clone sequences and assembly contigs with BWA-MEM. arXiv [Preprint] DOI: 10.48550/arXiv.1303.3997

Lukas D, Clutton-Brock TH. 2013. The evolution of social monogamy in mammals. Science. 341(6145):526–530.

Maretty L, Jensen JM, Petersen B, Sibbesen JA, Liu S, Villesen P, Skov L, Belling K, Theil Have C, Izarzugaza JMG, et al. 2017. Sequencing and *de novo* assembly of 150 genomes from Denmark as a population reference. Nature. 548(7665):87–91.

Mendel G. 1866. Versuche über Pflanzen-Hybriden. Verhandlungen des Naturforschenden Vereines, Abhandlungern, Brünn. 4:3–47.

Mendoza SP, Mason WA. 1986. Parental division of labour and differentiation of attachments in a monogamous primate (*Callicebus cupreus*). Anim Behav. 34(5):1336–1347.

Michaelson JJ, Shi Y, Gujral M, Zheng H, Malhotra D, Jin X, Jian M, Liu G, Greer D, Bhandari A, et al. 2012. Whole-genome sequencing in autism identifies hot spots for *de novo* germline mutation. Cell. 151(7):1431–1442.

Mohrenweiser HW, Wilson DM 3rd, Jones IM. 2003. Challenges and complexities in estimating both the functional impact and the disease risk associated with the extensive genetic variation in human DNA repair genes. Mutat Res. 526(1-2):93–125.

Nachman MW, Crowell SL. 2000. Estimate of the mutation rate per nucleotide in humans. Genetics. 156(1):297–304.

Pacifici M, Santini L, Di Marco M, Baisero D, Francucci L, Grottolo Marasini G, Visconti P, Rondinini C. 2013. Generation length for mammals. Nat Conserv. 5:87–94.

Patterson M, Marschall T, Pisanti N, van Iersel L, Stougie L, Klau GW, Schönhuth A. 2015. WhatsHap: weighted haplotype assembly for future-generation sequencing reads. J Comput Biol. 22(6):498–509.

Payer LM, Burns KH. 2019. Transposable elements in human genetic disease. Nat Rev Genet. 20(12):760–772.

Pfeifer SP. 2017a. Direct estimate of the spontaneous germ line mutation rate in African green monkeys. Evolution. 71(12):2858–2870.

Pfeifer SP. 2017b. From next-generation resequencing reads to a high-quality variant data set. Heredity (Edinb). 118(2):111–124.

Pfeifer SP. 2020. Spontaneous mutation rates. In: Ho SYW, editor. The molecular evolutionary clock. Theory and practice. Cham, Switzerland: Springer International Publishing; p. 35–44.

Pfeifer SP. 2021. Studying mutation rate evolution in primates-the effects of computational pipelines and parameter choices. GigaScience. 10(10):giab069.

Pfeifer SP, Baxter A, Savidge LE, Sedlazeck FJ, Bales KL. 2024. *De novo* genome assembly for the coppery titi monkey (*Plecturocebus cupreus*): an emerging nonhuman primate model for behavioral research. Genome Biol Evol. 16(5):evae108.

R Core Team. 2022. R: a language and environment for statistical computing. R Foundation for Statistical Computing, Vienna, Austria. https://www.R-project.org/.

Rahbari R, Wuster A, Lindsay SJ, Hardwick RJ, Alexandrov LB, Turki SA, Dominiczak A, Morris A, Porteous D, Smith B, et al. 2016. Timing, rates and spectra of human germline mutation. Nat Genet. 48(2):126–133.

Rigney N, de Vries GJ, Petrulis A, Young LJ. 2022. Oxytocin, vasopressin, and social behavior: from neural circuits to clinical opportunities. Endocrinology. 163(9):bqac111.

Sedlazeck FJ, Rescheneder P, Smolka M, Fang H, Nattestad M, von Haeseler A, Schatz MC. 2018. Accurate detection of complex structural variations using single-molecule sequencing. Nat Methods. 15(6):461–468.

Ségurel L, Wyman MJ, Przeworski M. 2014. Determinants of mutation rate variation in the human germline. Annu Rev Genomics Hum Genet.15(1):47–70.

Shendure J, Akey JM. 2015. The origins, determinants, and consequences of human mutations. Science. 349(6255):1478–1483.

Slotkin RK, Martienssen R. 2007. Transposable elements and the epigenetic regulation of the genome. Nat Rev Genet. 8(4):272–285.

Soni V, Pfeifer SP, Jensen JD. 2025c. Recent insights into the evolutionary genomics of the critically endangered aye-aye (*Daubentonia madagascariensis*). Am J Primatol. 87(12):e70105.

Soni V, Versoza C, Jensen JD, Pfeifer SP. 2025a. Inferring the landscape of mutation and recombination in the common marmoset (*Callithrix jacchus*) in the presence of twinning and hematopoietic chimerism. BioRxiv, preprint. DOI: 10.1101/2025.07.01.662565.

Soni V, Versoza C, Terbot J, Jensen JD, Pfeifer SP. 2025b. Inferring fine-scale mutation and recombination rate maps in ayes-ayes (*Daubentonia madagascariensis*). Ecol Evol. 15(11):e72314.

Spisak N, de Manuel M, Milligan W, Sella G, Przeworski M. 2024. The clock-like accumulation of germline and somatic mutations can arise from the interplay of DNA damage and repair. PLoS Biol. 22(6):e3002678.

Tatsumoto S, Go Y, Fukuta K, Noguchi H, Hayakawa T, Tomonaga M, Hirai H, Matsuzawa T, Agata K, Fujiyama A. 2017. Direct estimation of *de novo* mutation rates in a chimpanzee parent-offspring trio by ultra-deep whole genome sequencing. Sci Rep. 7(1):13561.

Thomas GWC, Wang RJ, Puri A, Harris RA, Raveendran M, Hughes DST, Murali SC, Williams LE, Doddapaneni H, Muzny DM, et al. 2018. Reproductive longevity predicts mutation rates in primates. Curr Biol. 28(19):3193–3197.

Thorvaldsdóttir H, Robinson JT, Mesirov JP. 2013. Integrative Genomics Viewer (IGV): high-performance genomics data visualization and exploration. Brief Bioinform. 14(2):178–192.

Tran LAP, Pfeifer SP. 2018. Germline mutation rates in Old World monkeys. Chichester: eLS, John Wiley & Sons, Ltd.

Valeggia CR, Mendoza SP, Fernandez-Duque E, Mason WA, Lasley B. 1999. Reproductive biology of female titi monkeys (*Callicebus moloch*) in captivity. Am J Primatol. 47(3):183–195.

Van Belle S, Fernandez-Duque E, Di Fiore A. 2016. Demography and life history of wild red titi monkeys (*Callicebus discolor*) and equatorial sakis (*Pithecia aequatorialis*) in Amazonian Ecuador: a 12-year study. Am J Primatol. 78(2):204–215.

van der Auwera GA, O’Connor BD. 2020. Genomics in the cloud: using Docker, GATK, and WDL in Terra. Sebastopol: O’Reilly Media.

Venn O, Turner I, Mathieson I, de Groot N, Bontrop R, McVean G. 2014. Strong male bias drives germline mutation in chimpanzees. Science. 344(6189):1272–1275.

Vermeer J, Baumeyer A. 2022. European *ex situ* programmes for larger New World monkeys. Neotrop Primates. 28(1-2).

Versoza CJ, Bales KL, Jensen JD, Pfeifer SP. 2026a. Sex-specific landscapes of crossover and non-crossover recombination in the coppery titi monkey (*Plecturocebus cupreus*). BioRxiv, preprint. DOI: 10.64898/2026.01.11.698870.

Versoza CJ, Bales KL, Jensen JD, Pfeifer SP. 2026b. The landscape of structural variation in coppery titi monkeys (*Plecturocebus cupreus*). BioRxiv, preprint.

Versoza CJ, Ehmke E, Jensen JD, Pfeifer SP. 2025. Characterizing the rates and patterns of *de novo* germline mutations in the aye-aye (*Daubentonia madagascariensis*). Mol Biol Evol. 42(3):msaf034.

Wang K, Li M, Hakonarson H. 2010. ANNOVAR: functional annotation of genetic variants from high-throughput sequencing data. Nucleic Acids Res. 38(16):e164.

Wang RJ, Thomas GWC, Raveendran M, Harris RA, Doddapaneni H, Muzny DM, Capitanio JP, Radivojac P, Rogers J, Hahn MW. 2020. Paternal age in rhesus macaques is positively associated with germline mutation accumulation but not with measures of offspring sociability. Genome Res. 30(6):826–834.

Wilson Sayres MA, Venditti C, Pagel M, Makova KD. 2011. Do variations in substitution rates and male mutation bias correlate with life-history traits? A study of 32 mammalian genomes. Evolution. 65(10):2800–2815.

Witczak LR, Samra J, Dufek M, Goetze LR, Freeman SM, Lau AR, Rothwell ES, Savidge LE, Arias-Del Razo R, Baxter A, et al. 2024. Expression of bond-related behaviors affects titi monkey responsiveness to oxytocin and vasopressin treatments. Ann NY Acad Sci. 1534(1):118–129.

Wong WS, Solomon BD, Bodian DL, Kothiyal P, Eley G, Huddleston KC, Baker R, Thach DC, Iyer RK, Vockley JG, et al. 2016. New observations on maternal age effect on germline *de novo* mutations. Nat Commun. 7:10486.

Wu FL, Strand AI, Cox LA, Ober C, Wall JD, Moorjani P, Przeworski M. 2020. A comparison of humans and baboons suggests germline mutation rates do not track cell divisions. PLoS Biol.18(8):e3000838.

Yang C, Zhou Y, Marcus S, Formenti G, Bergeron LA, Song Z, Bi X, Bergman J, Rousselle MMC, Zhou C, et al. 2021. Evolutionary and biomedical insights from a marmoset diploid genome assembly. Nature. 594(7862):227–233.

Zablocki-Thomas P, Lau A, Witczak L, Dufek M, Wright A, Savidge L, Paulus J, Baxter A, Karaskiewicz C, Seelke AMH, et al. 2023a. Intranasal oxytocin does not change partner preference in female titi monkeys (*Plecturocebus cupreus*), but intranasal vasopressin decreases it. J Neuroendocrinol. 35(10):e13339.

Zablocki-Thomas P, Rebout N, Karaskiewicz CL, Bales KL. 2023b. Survival rates and mortality risks of *Plecturocebus cupreus* at the California National Primate Research Center. Am J Primatol. 85(10):e23531.

Zhu T, Vats P, Onken S, Dunstan A, Zamirai B, Puleri DF, Nair A, Oliva M, Gaihre A, Sadhnani P, et al. 2025.Parabricks: GPU accelerated universal pan-instrument genomics analysis software suite. BioRxiv, preprint. DOI: 10.1101/2025.07.23.666378.

