## Supplementary Materials for "Characterizing the rates and patterns of *de novo* germline mutations in coppery titi monkeys (*Plecturocebus cupreus*)"

**Supplementary Table 1**. Sample information, including the ID and sex of the individuals, their parents' IDs and ages at the individual's birth, and mapping coverages.

| pedigree ID | sex | sire ID | dam ID | sire's age (in years) | dam's age (in years) | mapping coverage |
|---|---|---|---|---|---|---|
| 1 | M | – | – | – | – | 39.1 |
| 2 | F | – | – | – | – | 41.7 |
| 3 | M | – | – | – | – | 50.8 |
| 4 | F | – | – | – | – | 56.4 |
| 5 | M | – | – | – | – | 55.3 |
| 6 | F | – | – | – | – | 57.7 |
| 7 | M | – | – | – | – | 44.7 |
| 8 | F | 1 | 2 | 7.6 | 4.9 | 41.9 |
| 9 | M | 1 | 2 | 8.4 | 5.7 | 43.2 |
| 10 | F | – | – | – | – | 45.5 |
| 11 | F | 1 | 2 | 9.6 | 6.9 | 40.5 |
| 12 | F | 1 | 2 | 10.4 | 7.7 | 40.7 |
| 13 | M | – | – | – | – | 40.5 |
| 14 | M | 3 | 4 | 8.6 | 11.3 | 60.4 |
| 15 | M | 3 | 4 | 12.3 | 15.0 | 54.3 |
| 16 | F | – | – | – | – | 59.2 |
| 17 | M | 3 | 4 | 15.6 | 18.3 | 48.9 |
| 18 | F | 5 | 6 | 4.1 | 4.0 | 46.1 |
| 19 | M | 5 | 6 | 6.2 | 6.1 | 61.6 |
| 20 | M | 5 | 6 | 8.0 | 7.9 | 60.3 |
| 21 | M | 5 | 6 | 9.7 | 9.6 | 47.8 |
| 22 | F | 7 | 8 | 6.8 | 4.0 | 41.7 |
| 23 | M | 9 | 10 | 3.4 | 9.2 | 45.7 |
| 24 | F | 13 | 12 | 9.2 | 4.3 | 44.9 |
| 25 | F | 15 | 16 | 3.0 | 3.1 | 73.4 |

**Supplementary Table 2**. Summary of the autosomal single nucleotide polymorphism (SNP) data.

| chromosome | length | # SNPs | Ts/Tv |
|---|---|---|---|
| 1 | 235,103,535 | 1,771,735 | 2.54 |
| 2 | 158,738,102 | 1,167,278 | 2.57 |
| 3 | 165,937,890 | 1,275,720 | 2.47 |
| 4 | 151,623,469 | 1,046,581 | 2.55 |
| 5 | 150,993,727 | 1,160,479 | 2.50 |
| 6 | 123,740,199 | 875,613 | 2.57 |
| 7 | 135,512,294 | 1,071,622 | 2.44 |
| 8 | 116,834,941 | 890,566 | 2.40 |
| 9 | 101,234,152 | 792,897 | 2.50 |
| 10 | 75,878,286 | 592,687 | 2.68 |
| 11 | 49,373,254 | 383,620 | 2.58 |
| 12 | 231,176,762 | 1,787,064 | 2.54 |
| 13 | 118,029,634 | 786,396 | 2.65 |
| 14 | 130,776,705 | 858,039 | 2.57 |
| 15 | 95,533,141 | 692,513 | 2.51 |
| 16 | 94,172,006 | 657,758 | 2.49 |
| 17 | 73,180,778 | 544,597 | 2.73 |
| 18 | 92,880,398 | 788,665 | 2.44 |
| 19 | 65,768,754 | 507,354 | 2.80 |
| 20 | 91,752,637 | 696,754 | 2.36 |
| 21 | 73,432,007 | 538,439 | 2.45 |
| 22 | 44,717,611 | 310,780 | 2.45 |
| ∑ or ∅ | **2,576,390,282** | **19,197,157** | **2.54** |

**Supplementary Table 3**. Summary of the autosomal DNMs identified across the 15 parent-offspring trios. Locations (*chromosome [chr]* and *position*) and *genomic context* are defined according to the coppery titi monkey genome (GenBank accession number: GCA_040437455.1; Pfeifer et al. 2024). *REF* and *ALT* indicate the reference and alternate alleles, respectively. *Repeat status* indicates whether a DNM is located within a repetitive region of the genome. *CpG>TpG status* indicates C-to-T transitions at CpG sites. *Parent-of-origin* indicates the inferred parent-of-origin for a DNM (when available).

| chr | position | REF | ALT | recipient ID | transmitted? | parent-of-origin | repeat status | genomic context | CpG>TpG status |
|---|---|---|---|---|---|---|---|---|---|
| chr1 | 992,836 | C | T | 12 | YES | maternal | 0 | intergenic | 1 |
| chr1 | 1,138,179 | G | A | 24 | — | — | 1 | intergenic | 1 |
| chr1 | 2,494,351 | C | T | 24 | — | maternal | 0 | intergenic | 1 |
| chr1 | 3,710,073 | G | A | 14 | — | paternal | 1 | intergenic | 0 |
| chr1 | 14,014,473 | T | G | 18 | — | — | 0 | intergenic | 0 |
| chr1 | 31,707,269 | T | C | 19 | — | — | 1 | intronic | 0 |
| chr1 | 52,216,441 | C | T | 20 | — | — | 1 | intergenic | 1 |
| chr1 | 52,294,075 | A | C | 18 | — | paternal | 1 | intergenic | 0 |
| chr1 | 62,145,286 | C | T | 15 | NO | maternal | 1 | intronic | 0 |
| chr1 | 72,782,898 | C | A | 9 | NO | paternal | 0 | exonic | 0 |
| chr1 | 74,183,840 | C | T | 9 | NO | — | 1 | intergenic | 0 |
| chr1 | 76,888,030 | A | C | 20 | — | paternal | 1 | intronic | 0 |
| chr1 | 78,279,184 | A | T | 15 | YES | paternal | 1 | intergenic | 0 |
| chr1 | 80,337,778 | G | A | 8 | NO | paternal | 0 | intergenic | 0 |
| chr1 | 82,439,804 | T | C | 15 | YES | paternal | 1 | intergenic | 0 |
| chr1 | 83,567,449 | C | T | 8 | YES | paternal | 0 | intergenic | 1 |
| chr1 | 93,974,165 | T | G | 25 | — | — | 0 | intergenic | 0 |
| chr1 | 101,859,185 | T | C | 22 | — | maternal | 0 | intergenic | 0 |
| chr1 | 102,802,695 | T | G | 8 | YES | paternal | 1 | intergenic | 0 |
| chr1 | 105,854,375 | T | G | 8 | NO | — | 0 | intronic | 0 |
| chr1 | 114,045,875 | C | T | 8 | YES | paternal | 0 | intergenic | 0 |
| chr1 | 115,976,408 | A | T | 23 | — | maternal | 0 | intergenic | 0 |
| chr1 | 117,548,528 | T | C | 23 | — | — | 0 | intergenic | 0 |
| chr1 | 123,384,408 | A | C | 25 | — | paternal | 0 | intergenic | 0 |
| chr1 | 126,520,125 | G | A | 9 | YES | — | 1 | intronic | 0 |
| chr1 | 130,182,960 | C | T | 23 | — | paternal | 1 | upstream | 1 |
| chr1 | 131,074,499 | G | A | 23 | — | — | 1 | intergenic | 1 |
| chr1 | 131,850,253 | C | T | 8 | NO | — | 0 | intergenic | 0 |
| chr1 | 134,860,365 | G | A | 22 | — | — | 1 | intergenic | 0 |
| chr1 | 143,200,186 | C | T | 22 | — | paternal | 1 | intronic | 0 |
| chr1 | 148,762,350 | C | T | 11 | — | paternal | 1 | intergenic | 0 |
| chr1 | 154,968,109 | G | A | 11 | — | paternal | 0 | intronic | 0 |
| chr1 | 156,766,674 | G | A | 8 | NO | maternal | 1 | intergenic | 1 |
| chr1 | 158,536,783 | A | G | 12 | YES | paternal | 1 | intergenic | 0 |
| chr1 | 159,817,330 | G | A | 19 | — | — | 1 | intergenic | 1 |
| chr1 | 160,025,237 | G | C | 23 | — | — | 0 | intergenic | 0 |
| chr1 | 160,262,364 | C | A | 24 | — | paternal | 0 | intergenic | 0 |
| chr1 | 165,775,094 | C | T | 8 | YES | paternal | 0 | intronic | 0 |
| chr1 | 173,631,154 | G | A | 23 | — | paternal | 1 | intergenic | 0 |
| chr1 | 174,984,067 | C | G | 20 | — | — | 0 | intergenic | 0 |
| chr1 | 176,736,933 | G | C | 8 | YES | — | 1 | intergenic | 0 |
| chr1 | 180,002,068 | G | A | 14 | — | — | 0 | intergenic | 0 |
| chr1 | 180,442,376 | G | A | 14 | — | — | 0 | intergenic | 1 |
| chr1 | 181,343,066 | A | G | 9 | NO | paternal | 0 | intergenic | 0 |
| chr1 | 191,688,707 | A | C | 12 | NO | — | 1 | intergenic | 0 |
| chr1 | 195,566,203 | T | G | 21 | — | paternal | 0 | intergenic | 0 |
| chr1 | 196,777,588 | C | T | 15 | YES | paternal | 1 | intergenic | 1 |
| chr1 | 196,819,805 | G | A | 22 | — | maternal | 1 | intergenic | 1 |
| chr1 | 207,525,039 | G | C | 23 | — | paternal | 1 | intergenic | 0 |

| | | | | | | | | | |
|---|---|---|---|---|---|---|---|---|---|
| chr1 | 208,911,753 | A | T | 21 | — | paternal | 0 | intergenic | 0 |
| chr1 | 214,592,146 | G | T | 25 | — | maternal | 0 | intergenic | 0 |
| chr1 | 219,756,174 | A | G | 23 | — | paternal | 0 | intergenic | 0 |
| chr1 | 223,462,636 | C | A | 23 | — | paternal | 0 | intergenic | 0 |
| chr1 | 230,855,509 | G | A | 23 | — | paternal | 0 | intergenic | 0 |
| chr1 | 233,267,746 | A | C | 8 | YES | paternal | 1 | intergenic | 0 |
| chr2 | 15,828,710 | A | G | 15 | YES | paternal | 1 | intergenic | 0 |
| chr2 | 32,734,618 | C | A | 23 | — | paternal | 1 | intronic | 0 |
| chr2 | 33,343,920 | C | T | 23 | — | maternal | 0 | intergenic | 0 |
| chr2 | 33,384,557 | A | G | 18 | — | — | 0 | intergenic | 0 |
| chr2 | 40,905,385 | G | C | 21 | — | paternal | 0 | intergenic | 0 |
| chr2 | 46,339,685 | A | C | 23 | — | maternal | 0 | exonic | 0 |
| chr2 | 47,215,311 | C | T | 23 | — | — | 0 | intergenic | 0 |
| chr2 | 50,958,560 | A | G | 24 | — | — | 0 | intergenic | 0 |
| chr2 | 54,963,871 | C | T | 20 | — | paternal | 1 | intergenic | 1 |
| chr2 | 55,503,240 | T | C | 14 | — | paternal | 0 | intergenic | 0 |
| chr2 | 64,757,046 | G | A | 14 | — | — | 0 | intergenic | 0 |
| chr2 | 71,887,814 | T | A | 24 | — | — | 1 | intergenic | 0 |
| chr2 | 79,394,340 | A | C | 24 | — | maternal | 0 | intergenic | 0 |
| chr2 | 93,773,009 | G | A | 12 | NO | — | 0 | intergenic | 0 |
| chr2 | 97,187,713 | A | G | 14 | — | — | 0 | intergenic | 0 |
| chr2 | 98,758,158 | A | G | 19 | — | — | 1 | intergenic | 0 |
| chr2 | 100,055,487 | C | G | 15 | YES | paternal | 0 | intergenic | 0 |
| chr2 | 104,150,649 | G | T | 11 | — | maternal | 1 | intergenic | 0 |
| chr2 | 107,559,785 | C | T | 9 | YES | maternal | 0 | intergenic | 0 |
| chr2 | 121,707,200 | C | T | 11 | — | — | 1 | intergenic | 1 |
| chr2 | 132,181,026 | G | A | 18 | — | — | 0 | intergenic | 1 |
| chr2 | 152,872,455 | G | A | 8 | NO | — | 1 | intergenic | 1 |
| chr3 | 738,481 | G | A | 23 | — | maternal | 0 | intergenic | 0 |
| chr3 | 6,532,416 | T | C | 23 | — | paternal | 1 | intergenic | 0 |
| chr3 | 8,755,583 | T | G | 21 | — | — | 0 | intronic | 0 |
| chr3 | 14,743,162 | G | A | 15 | NO | paternal | 0 | intergenic | 0 |
| chr3 | 16,620,542 | A | G | 22 | — | maternal | 1 | intergenic | 0 |
| chr3 | 25,915,206 | A | G | 23 | — | — | 0 | intronic | 0 |
| chr3 | 32,491,318 | A | C | 9 | NO | paternal | 0 | intergenic | 0 |
| chr3 | 41,702,909 | A | C | 18 | — | paternal | 1 | intergenic | 0 |
| chr3 | 45,055,131 | A | C | 15 | NO | paternal | 0 | intergenic | 0 |
| chr3 | 47,251,529 | C | T | 18 | — | paternal | 0 | intergenic | 1 |
| chr3 | 54,029,715 | T | C | 15 | NO | paternal | 1 | intronic | 0 |
| chr3 | 56,924,868 | G | A | 25 | — | paternal | 1 | intergenic | 1 |
| chr3 | 74,220,414 | T | C | 19 | — | paternal | 1 | intergenic | 0 |
| chr3 | 74,359,543 | C | T | 12 | NO | paternal | 1 | intergenic | 1 |
| chr3 | 77,949,571 | G | A | 20 | — | paternal | 1 | intergenic | 0 |
| chr3 | 81,131,490 | T | G | 14 | — | paternal | 1 | intergenic | 0 |
| chr3 | 82,289,338 | T | A | 25 | — | — | 0 | intergenic | 0 |
| chr3 | 87,584,912 | C | G | 23 | — | paternal | 1 | intergenic | 0 |
| chr3 | 89,118,313 | G | C | 15 | NO | paternal | 1 | intergenic | 0 |
| chr3 | 99,125,508 | G | A | 20 | — | paternal | 0 | intergenic | 0 |
| chr3 | 103,368,687 | C | T | 8 | NO | maternal | 1 | intergenic | 0 |
| chr3 | 116,341,622 | C | G | 19 | — | — | 1 | intergenic | 0 |
| chr3 | 116,655,794 | A | G | 21 | — | — | 0 | intergenic | 0 |
| chr3 | 117,972,530 | G | A | 14 | — | — | 0 | intergenic | 1 |
| chr3 | 124,809,977 | T | G | 18 | — | maternal | 0 | intergenic | 0 |
| chr3 | 127,711,966 | C | T | 12 | NO | — | 1 | intergenic | 1 |
| chr3 | 142,994,741 | T | C | 23 | — | maternal | 1 | intronic | 0 |
| chr3 | 143,580,671 | G | A | 22 | — | paternal | 0 | intergenic | 1 |
| chr3 | 146,860,931 | C | T | 12 | YES | paternal | 0 | intergenic | 1 |
| chr3 | 155,051,909 | G | A | 18 | — | — | 0 | intergenic | 1 |
| chr3 | 156,640,792 | C | T | 15 | YES | paternal | 0 | intronic | 1 |

| | | | | | | | | | |
|---|---|---|---|---|---|---|---|---|---|
| chr3 | 157,882,350 | G | T | 25 | — | maternal | 0 | intergenic | 0 |
| chr3 | 161,176,593 | A | G | 9 | NO | — | 0 | intronic | 0 |
| chr4 | 867,918 | G | A | 19 | — | paternal | 0 | intergenic | 0 |
| chr4 | 10,777,640 | G | A | 20 | — | — | 0 | intergenic | 0 |
| chr4 | 20,085,806 | A | G | 23 | — | — | 0 | intergenic | 0 |
| chr4 | 20,765,625 | C | T | 25 | — | paternal | 1 | intergenic | 1 |
| chr4 | 30,543,869 | G | A | 15 | NO | — | 1 | exonic | 0 |
| chr4 | 37,501,523 | G | A | 23 | — | paternal | 0 | exonic | 1 |
| chr4 | 41,651,010 | A | G | 11 | — | — | 0 | upstream | 0 |
| chr4 | 45,009,645 | G | T | 19 | — | paternal | 0 | intergenic | 0 |
| chr4 | 46,552,217 | A | C | 12 | NO | — | 0 | exonic | 0 |
| chr4 | 65,630,681 | G | A | 12 | NO | paternal | 0 | intergenic | 0 |
| chr4 | 69,086,744 | C | T | 24 | — | maternal | 0 | intronic | 0 |
| chr4 | 73,811,688 | G | T | 23 | — | maternal | 0 | intergenic | 0 |
| chr4 | 87,056,880 | T | G | 15 | NO | paternal | 1 | intergenic | 0 |
| chr4 | 87,614,003 | G | A | 11 | — | maternal | 0 | intergenic | 1 |
| chr4 | 93,701,643 | C | G | 24 | — | paternal | 0 | intergenic | 0 |
| chr4 | 97,711,016 | G | C | 23 | — | — | 0 | intergenic | 0 |
| chr4 | 101,605,440 | T | C | 15 | YES | — | 0 | intergenic | 0 |
| chr4 | 108,910,289 | A | C | 25 | — | paternal | 0 | intergenic | 0 |
| chr4 | 111,190,140 | T | G | 18 | — | — | 0 | intergenic | 0 |
| chr4 | 112,988,082 | T | C | 25 | — | maternal | 0 | intronic | 0 |
| chr4 | 132,481,456 | C | T | 11 | — | — | 0 | intronic | 0 |
| chr4 | 136,860,793 | C | T | 23 | — | — | 0 | downstream | 0 |
| chr4 | 143,256,228 | T | G | 11 | — | paternal | 1 | intergenic | 0 |
| chr4 | 146,354,213 | G | A | 18 | — | — | 0 | intergenic | 1 |
| chr5 | 5,567,090 | G | A | 18 | — | — | 0 | intergenic | 1 |
| chr5 | 7,865,258 | G | A | 8 | YES | paternal | 1 | intergenic | 0 |
| chr5 | 18,671,569 | A | T | 21 | — | paternal | 1 | intergenic | 0 |
| chr5 | 20,431,561 | T | C | 24 | — | paternal | 0 | intergenic | 0 |
| chr5 | 24,056,211 | G | A | 12 | NO | maternal | 0 | intergenic | 0 |
| chr5 | 31,145,366 | G | A | 24 | — | — | 0 | intergenic | 0 |
| chr5 | 39,677,229 | A | G | 20 | — | — | 0 | intergenic | 0 |
| chr5 | 43,216,886 | G | A | 19 | — | — | 1 | intergenic | 1 |
| chr5 | 50,205,466 | C | T | 8 | YES | paternal | 0 | intergenic | 1 |
| chr5 | 51,872,911 | C | T | 11 | — | paternal | 0 | intergenic | 1 |
| chr5 | 52,513,173 | G | A | 19 | — | maternal | 1 | intergenic | 0 |
| chr5 | 55,517,954 | T | G | 8 | NO | — | 1 | intronic | 0 |
| chr5 | 60,263,937 | C | T | 18 | — | maternal | 0 | intronic | 0 |
| chr5 | 86,091,407 | G | A | 18 | — | maternal | 0 | intergenic | 0 |
| chr5 | 86,481,748 | G | A | 8 | YES | paternal | 0 | intronic | 0 |
| chr5 | 94,033,883 | C | T | 9 | NO | paternal | 0 | intergenic | 0 |
| chr5 | 96,084,926 | C | T | 23 | — | paternal | 0 | intronic | 0 |
| chr5 | 109,971,378 | C | T | 23 | — | — | 0 | intergenic | 0 |
| chr5 | 115,827,434 | T | G | 18 | — | paternal | 0 | intergenic | 0 |
| chr5 | 121,262,825 | C | A | 23 | — | — | 1 | intergenic | 0 |
| chr5 | 124,683,851 | A | C | 23 | — | — | 0 | intronic | 0 |
| chr5 | 130,200,728 | A | G | 24 | — | paternal | 0 | intergenic | 0 |
| chr5 | 133,940,796 | G | A | 14 | — | paternal | 0 | intergenic | 1 |
| chr5 | 136,616,213 | C | T | 23 | — | maternal | 1 | intronic | 0 |
| chr5 | 137,387,324 | C | A | 18 | — | paternal | 1 | intergenic | 0 |
| chr5 | 145,088,661 | T | C | 8 | NO | — | 1 | intergenic | 0 |
| chr6 | 4,140,849 | C | T | 25 | — | paternal | 0 | intergenic | 0 |
| chr6 | 7,952,237 | C | T | 21 | — | paternal | 1 | intergenic | 1 |
| chr6 | 13,509,272 | T | G | 9 | NO | paternal | 1 | intergenic | 0 |
| chr6 | 18,878,363 | C | T | 9 | YES | paternal | 0 | intergenic | 0 |
| chr6 | 24,822,543 | C | G | 8 | YES | paternal | 0 | intergenic | 0 |
| chr6 | 25,969,320 | G | A | 18 | — | paternal | 0 | exonic | 1 |
| chr6 | 35,551,448 | G | A | 9 | YES | paternal | 0 | intergenic | 1 |

| | | | | | | | | | |
|---|---|---|---|---|---|---|---|---|---|
| chr6 | 44,071,714 | C | T | 25 | — | paternal | 0 | intronic | 0 |
| chr6 | 50,393,447 | A | T | 19 | — | paternal | 0 | intronic | 0 |
| chr6 | 56,299,590 | C | T | 12 | NO | paternal | 0 | intergenic | 0 |
| chr6 | 62,572,831 | C | T | 23 | — | paternal | 1 | intergenic | 1 |
| chr6 | 63,577,808 | G | C | 20 | — | paternal | 0 | intergenic | 0 |
| chr6 | 63,610,479 | C | G | 14 | — | paternal | 1 | intergenic | 0 |
| chr6 | 65,998,693 | A | C | 11 | — | — | 0 | intergenic | 0 |
| chr6 | 69,492,936 | C | T | 21 | — | paternal | 0 | intergenic | 0 |
| chr6 | 70,718,857 | C | T | 23 | — | paternal | 1 | intergenic | 0 |
| chr6 | 75,107,584 | G | T | 8 | NO | paternal | 0 | intergenic | 0 |
| chr6 | 75,659,732 | G | A | 14 | — | paternal | 1 | downstream | 1 |
| chr6 | 76,773,216 | G | A | 8 | NO | paternal | 1 | intronic | 1 |
| chr6 | 78,553,861 | G | A | 8 | NO | paternal | 1 | intergenic | 1 |
| chr6 | 91,144,110 | G | T | 12 | NO | paternal | 1 | intergenic | 0 |
| chr6 | 92,406,406 | C | T | 15 | NO | paternal | 1 | exonic | 1 |
| chr6 | 94,424,476 | A | G | 9 | YES | — | 0 | intergenic | 0 |
| chr6 | 96,439,012 | C | G | 25 | — | paternal | 0 | intronic | 0 |
| chr6 | 97,203,640 | C | A | 20 | — | — | 1 | intergenic | 0 |
| chr6 | 108,942,230 | G | A | 15 | YES | paternal | 1 | intergenic | 0 |
| chr6 | 116,615,790 | C | T | 23 | — | — | 1 | intergenic | 0 |
| chr7 | 8,047,038 | G | A | 21 | — | paternal | 0 | intergenic | 0 |
| chr7 | 8,741,012 | A | G | 21 | — | maternal | 1 | intronic | 0 |
| chr7 | 15,404,580 | G | T | 18 | — | maternal | 0 | intergenic | 0 |
| chr7 | 18,219,947 | C | T | 22 | — | — | 0 | intergenic | 0 |
| chr7 | 24,856,795 | T | C | 9 | NO | — | 0 | intergenic | 0 |
| chr7 | 44,865,518 | A | C | 11 | — | paternal | 0 | intergenic | 0 |
| chr7 | 54,414,570 | C | T | 19 | — | — | 0 | intronic | 0 |
| chr7 | 56,736,412 | A | C | 9 | NO | paternal | 1 | intronic | 0 |
| chr7 | 66,511,484 | A | C | 25 | — | paternal | 1 | intronic | 0 |
| chr7 | 67,982,451 | T | C | 23 | — | paternal | 1 | intergenic | 0 |
| chr7 | 68,845,484 | A | G | 23 | — | — | 1 | intergenic | 0 |
| chr7 | 72,673,987 | G | A | 25 | — | maternal | 0 | intergenic | 1 |
| chr7 | 83,027,713 | G | A | 8 | NO | — | 1 | intronic | 1 |
| chr7 | 93,169,750 | T | C | 14 | — | maternal | 0 | intergenic | 0 |
| chr7 | 94,197,623 | C | T | 23 | — | paternal | 0 | upstream | 1 |
| chr7 | 97,443,615 | G | A | 14 | — | paternal | 0 | intergenic | 0 |
| chr7 | 104,292,894 | T | C | 22 | — | — | 0 | intergenic | 0 |
| chr7 | 121,443,817 | T | C | 19 | — | — | 1 | intergenic | 0 |
| chr7 | 131,292,573 | A | T | 22 | — | paternal | 0 | intergenic | 0 |
| chr8 | 475,833 | C | G | 19 | — | paternal | 1 | intergenic | 0 |
| chr8 | 1,397,861 | C | T | 14 | — | — | 1 | intergenic | 1 |
| chr8 | 4,001,245 | A | G | 23 | — | — | 0 | intergenic | 0 |
| chr8 | 4,080,951 | G | A | 15 | YES | paternal | 1 | intergenic | 0 |
| chr8 | 7,721,361 | A | C | 25 | — | paternal | 0 | intronic | 0 |
| chr8 | 10,400,414 | G | A | 15 | YES | paternal | 0 | intergenic | 0 |
| chr8 | 35,598,691 | C | T | 11 | — | — | 0 | intergenic | 0 |
| chr8 | 39,388,594 | G | A | 24 | — | — | 1 | intergenic | 1 |
| chr8 | 60,521,900 | A | G | 24 | — | paternal | 1 | upstream | 0 |
| chr8 | 67,900,825 | A | C | 20 | — | paternal | 1 | intergenic | 0 |
| chr8 | 72,580,912 | T | A | 12 | YES | — | 0 | intergenic | 0 |
| chr8 | 81,122,907 | G | A | 21 | — | paternal | 0 | intergenic | 0 |
| chr8 | 91,014,650 | T | A | 15 | NO | paternal | 0 | intergenic | 0 |
| chr8 | 91,058,077 | C | T | 20 | — | — | 0 | intergenic | 0 |
| chr8 | 93,517,104 | T | C | 22 | — | — | 0 | intergenic | 0 |
| chr8 | 109,049,991 | G | A | 18 | — | paternal | 0 | intergenic | 1 |
| chr8 | 109,636,896 | C | T | 14 | — | paternal | 1 | intergenic | 1 |
| chr8 | 113,052,633 | G | A | 11 | — | — | 1 | intronic | 0 |
| chr9 | 1,784,255 | G | A | 14 | — | paternal | 0 | intronic | 1 |
| chr9 | 6,570,439 | A | G | 15 | YES | paternal | 0 | intronic | 0 |

| | | | | | | | | | |
|---|---|---|---|---|---|---|---|---|---|
| chr9 | 17,332,862 | C | T | 8 | NO | maternal | 1 | intergenic | 1 |
| chr9 | 21,652,064 | G | T | 22 | — | maternal | 0 | intergenic | 0 |
| chr9 | 25,000,912 | C | T | 8 | YES | paternal | 1 | intergenic | 0 |
| chr9 | 27,550,654 | T | C | 24 | — | paternal | 0 | intronic | 0 |
| chr9 | 38,684,888 | C | T | 15 | YES | — | 1 | intergenic | 0 |
| chr9 | 39,327,688 | G | A | 25 | — | paternal | 1 | intergenic | 0 |
| chr9 | 98,290,182 | G | A | 19 | — | paternal | 1 | intergenic | 1 |
| chr9 | 100,290,949 | A | C | 20 | — | — | 0 | intergenic | 0 |
| chr10 | 4,478,113 | T | G | 20 | — | — | 0 | intergenic | 0 |
| chr10 | 9,520,353 | C | G | 9 | NO | — | 0 | intergenic | 0 |
| chr10 | 19,840,340 | G | A | 15 | YES | — | 0 | downstream | 1 |
| chr10 | 21,293,246 | G | A | 20 | — | — | 1 | downstream | 0 |
| chr10 | 24,351,960 | T | G | 9 | NO | paternal | 1 | intergenic | 0 |
| chr10 | 26,326,691 | C | G | 23 | — | paternal | 0 | intronic | 0 |
| chr10 | 27,291,785 | T | C | 23 | — | — | 1 | intergenic | 0 |
| chr10 | 29,005,756 | C | T | 9 | YES | paternal | 0 | intronic | 0 |
| chr10 | 39,554,695 | C | T | 24 | — | paternal | 0 | intergenic | 1 |
| chr10 | 55,083,439 | C | T | 24 | — | paternal | 0 | upstream | 1 |
| chr10 | 57,286,034 | C | A | 8 | NO | — | 0 | intergenic | 0 |
| chr10 | 60,279,121 | G | A | 20 | — | maternal | 1 | intronic | 0 |
| chr11 | 4,404,554 | A | G | 11 | — | paternal | 1 | intergenic | 0 |
| chr11 | 4,414,546 | A | T | 12 | NO | maternal | 0 | intergenic | 0 |
| chr11 | 4,875,321 | T | A | 21 | — | — | 0 | intergenic | 0 |
| chr11 | 16,284,176 | G | C | 22 | — | paternal | 0 | intergenic | 0 |
| chr11 | 26,877,186 | G | C | 24 | — | paternal | 0 | intergenic | 0 |
| chr11 | 39,907,815 | C | T | 23 | — | — | 0 | intergenic | 0 |
| chr11 | 42,592,912 | C | A | 23 | — | paternal | 1 | intergenic | 0 |
| chr12 | 9,580,236 | C | T | 19 | — | — | 0 | intergenic | 0 |
| chr12 | 10,972,174 | G | A | 19 | — | — | 1 | intergenic | 0 |
| chr12 | 27,491,874 | A | T | 23 | — | paternal | 1 | intergenic | 0 |
| chr12 | 31,233,739 | C | T | 21 | — | paternal | 0 | intergenic | 0 |
| chr12 | 43,007,948 | C | T | 22 | — | — | 1 | intergenic | 1 |
| chr12 | 51,271,656 | C | G | 20 | — | — | 0 | intergenic | 0 |
| chr12 | 54,077,021 | G | A | 20 | — | paternal | 1 | intergenic | 0 |
| chr12 | 55,237,949 | T | A | 23 | — | — | 0 | intronic | 0 |
| chr12 | 68,430,060 | T | C | 19 | — | — | 0 | exonic | 0 |
| chr12 | 74,649,638 | G | A | 15 | YES | paternal | 0 | downstream | 1 |
| chr12 | 83,502,209 | T | C | 15 | NO | maternal | 0 | intergenic | 0 |
| chr12 | 93,662,581 | A | C | 21 | — | maternal | 1 | intergenic | 0 |
| chr12 | 114,466,569 | C | T | 8 | YES | — | 0 | intronic | 0 |
| chr12 | 114,643,474 | G | A | 12 | YES | paternal | 0 | downstream | 0 |
| chr12 | 125,709,830 | G | C | 23 | — | — | 0 | intergenic | 0 |
| chr12 | 135,558,640 | G | A | 11 | — | paternal | 1 | intergenic | 0 |
| chr12 | 135,580,356 | C | T | 8 | YES | — | 1 | intergenic | 0 |
| chr12 | 154,086,546 | T | C | 22 | — | — | 0 | exonic | 0 |
| chr12 | 160,500,241 | G | A | 9 | YES | maternal | 0 | upstream | 0 |
| chr12 | 163,467,203 | T | G | 15 | YES | — | 0 | intronic | 0 |
| chr12 | 165,395,724 | C | T | 14 | — | paternal | 0 | intronic | 1 |
| chr12 | 165,635,483 | C | T | 22 | — | — | 0 | intergenic | 0 |
| chr12 | 165,689,467 | G | A | 11 | — | maternal | 1 | intergenic | 1 |
| chr12 | 168,350,791 | G | A | 19 | — | paternal | 0 | intronic | 1 |
| chr12 | 177,084,013 | C | T | 18 | — | paternal | 1 | intronic | 0 |
| chr12 | 179,377,327 | C | T | 12 | YES | paternal | 0 | intergenic | 1 |
| chr12 | 182,932,344 | A | G | 21 | — | maternal | 1 | intergenic | 0 |
| chr12 | 185,488,288 | G | C | 23 | — | maternal | 0 | intergenic | 0 |
| chr12 | 191,514,077 | G | A | 12 | YES | paternal | 0 | upstream | 1 |
| chr12 | 199,415,735 | A | C | 21 | — | paternal | 0 | intronic | 0 |
| chr12 | 206,727,662 | C | T | 15 | NO | paternal | 0 | intergenic | 1 |
| chr12 | 212,987,478 | C | T | 25 | — | maternal | 1 | intergenic | 0 |

| | | | | | | | | | |
|---|---|---|---|---|---|---|---|---|---|
| chr12 | 223,157,855 | C | G | 12 | NO | — | 0 | intergenic | 0 |
| chr12 | 224,808,532 | T | G | 23 | — | maternal | 0 | intergenic | 0 |
| chr12 | 225,448,748 | G | C | 23 | — | maternal | 0 | intergenic | 0 |
| chr13 | 3,197,662 | G | A | 19 | — | paternal | 0 | intronic | 0 |
| chr13 | 21,226,898 | C | T | 9 | YES | paternal | 0 | intergenic | 0 |
| chr13 | 45,869,243 | G | A | 21 | — | paternal | 0 | intergenic | 1 |
| chr13 | 53,047,037 | A | C | 12 | YES | paternal | 1 | intronic | 0 |
| chr13 | 69,413,105 | T | C | 20 | — | — | 0 | intergenic | 0 |
| chr13 | 73,059,260 | T | C | 21 | — | — | 0 | intergenic | 0 |
| chr13 | 79,964,933 | C | A | 14 | — | — | 1 | intergenic | 0 |
| chr13 | 80,943,050 | A | C | 11 | — | — | 0 | intronic | 0 |
| chr13 | 81,784,412 | C | T | 23 | — | paternal | 0 | intergenic | 1 |
| chr13 | 84,889,926 | G | A | 8 | NO | — | 0 | intergenic | 0 |
| chr13 | 89,549,015 | C | T | 8 | NO | — | 1 | intergenic | 0 |
| chr13 | 90,794,411 | G | A | 19 | — | maternal | 1 | intergenic | 0 |
| chr13 | 96,262,105 | C | T | 15 | NO | paternal | 1 | intronic | 0 |
| chr13 | 102,174,428 | C | T | 23 | — | paternal | 0 | intergenic | 0 |
| chr13 | 108,120,172 | G | A | 20 | — | — | 1 | exonic | 0 |
| chr13 | 109,570,750 | C | T | 12 | NO | — | 0 | intronic | 0 |
| chr14 | 7,171,304 | G | C | 19 | — | paternal | 1 | intergenic | 0 |
| chr14 | 8,711,780 | A | G | 23 | — | paternal | 0 | intergenic | 0 |
| chr14 | 14,010,964 | C | T | 18 | — | paternal | 0 | intergenic | 0 |
| chr14 | 15,544,214 | C | T | 23 | — | paternal | 0 | intronic | 0 |
| chr14 | 20,995,811 | G | T | 23 | — | paternal | 0 | intergenic | 0 |
| chr14 | 25,179,129 | A | C | 19 | — | paternal | 1 | intergenic | 0 |
| chr14 | 34,149,243 | A | C | 18 | — | paternal | 1 | exonic | 0 |
| chr14 | 44,200,614 | C | T | 25 | — | — | 1 | intronic | 0 |
| chr14 | 44,846,807 | G | A | 22 | — | — | 1 | intergenic | 0 |
| chr14 | 46,351,965 | C | T | 15 | YES | paternal | 0 | upstream | 1 |
| chr14 | 47,061,354 | A | T | 8 | NO | paternal | 1 | intronic | 0 |
| chr14 | 54,341,708 | G | A | 21 | — | paternal | 0 | intergenic | 0 |
| chr14 | 64,845,693 | C | T | 22 | — | paternal | 0 | intergenic | 0 |
| chr14 | 71,703,577 | G | A | 18 | — | paternal | 1 | intergenic | 0 |
| chr14 | 71,932,807 | G | A | 19 | — | — | 1 | intronic | 1 |
| chr14 | 72,211,489 | A | G | 23 | — | — | 0 | intergenic | 0 |
| chr14 | 80,744,590 | G | A | 21 | — | paternal | 0 | intronic | 0 |
| chr14 | 87,290,545 | T | G | 9 | NO | — | 0 | intergenic | 0 |
| chr14 | 90,176,839 | A | T | 14 | — | paternal | 1 | intronic | 0 |
| chr14 | 92,727,436 | C | G | 23 | — | paternal | 0 | intronic | 0 |
| chr14 | 93,447,036 | C | T | 15 | YES | maternal | 0 | exonic | 0 |
| chr14 | 105,250,209 | C | A | 23 | — | paternal | 1 | intergenic | 0 |
| chr14 | 105,260,326 | G | C | 23 | — | — | 0 | intergenic | 0 |
| chr14 | 108,848,950 | C | T | 19 | — | — | 0 | intergenic | 1 |
| chr14 | 109,081,166 | C | A | 15 | NO | — | 0 | intergenic | 0 |
| chr15 | 4,465,513 | T | C | 8 | YES | — | 0 | intronic | 0 |
| chr15 | 8,609,643 | G | A | 14 | — | maternal | 0 | upstream | 0 |
| chr15 | 10,006,731 | T | C | 21 | — | paternal | 0 | intergenic | 0 |
| chr15 | 16,927,624 | C | T | 14 | — | paternal | 1 | intergenic | 0 |
| chr15 | 22,058,096 | C | T | 8 | NO | maternal | 0 | intronic | 0 |
| chr15 | 27,929,304 | G | C | 18 | — | paternal | 1 | intergenic | 0 |
| chr15 | 41,249,621 | C | T | 14 | — | paternal | 0 | intergenic | 0 |
| chr15 | 55,444,636 | A | C | 20 | — | — | 1 | intergenic | 0 |
| chr15 | 56,842,290 | C | T | 22 | — | paternal | 1 | intergenic | 0 |
| chr15 | 57,738,231 | A | T | 14 | — | — | 0 | intergenic | 0 |
| chr15 | 58,136,440 | T | G | 18 | — | maternal | 1 | intergenic | 0 |
| chr15 | 68,725,324 | G | T | 24 | — | — | 0 | intergenic | 0 |
| chr15 | 78,771,216 | G | C | 21 | — | paternal | 1 | intergenic | 0 |
| chr15 | 80,256,849 | G | T | 21 | — | paternal | 0 | intergenic | 0 |
| chr15 | 83,961,499 | A | C | 23 | — | — | 1 | intergenic | 0 |

| | | | | | | | | | |
|---|---|---|---|---|---|---|---|---|---|
| chr15 | 84,418,771 | G | A | 21 | — | paternal | 0 | intergenic | 1 |
| chr15 | 91,669,198 | T | C | 24 | — | maternal | 1 | intergenic | 0 |
| chr16 | 2,726,577 | C | T | 9 | NO | paternal | 1 | intergenic | 1 |
| chr16 | 3,391,053 | G | A | 11 | — | paternal | 0 | intergenic | 1 |
| chr16 | 3,816,034 | C | T | 15 | YES | paternal | 0 | intergenic | 0 |
| chr16 | 8,868,699 | G | A | 19 | — | paternal | 1 | intergenic | 1 |
| chr16 | 12,108,227 | G | A | 9 | YES | — | 1 | intergenic | 0 |
| chr16 | 12,912,027 | A | C | 12 | NO | paternal | 0 | intergenic | 0 |
| chr16 | 13,565,067 | T | G | 18 | — | paternal | 1 | intergenic | 0 |
| chr16 | 15,367,568 | A | C | 11 | — | maternal | 0 | intergenic | 0 |
| chr16 | 20,994,085 | C | T | 15 | YES | — | 1 | intergenic | 1 |
| chr16 | 22,101,749 | T | C | 18 | — | maternal | 0 | intronic | 0 |
| chr16 | 25,332,074 | A | C | 8 | NO | — | 0 | intergenic | 0 |
| chr16 | 29,559,525 | C | T | 23 | — | maternal | 1 | intergenic | 0 |
| chr16 | 32,280,785 | C | T | 19 | — | paternal | 0 | intergenic | 1 |
| chr16 | 38,387,516 | A | G | 25 | — | paternal | 1 | intronic | 0 |
| chr16 | 48,518,377 | T | G | 23 | — | paternal | 0 | intronic | 0 |
| chr16 | 51,303,589 | A | T | 15 | NO | — | 1 | intergenic | 0 |
| chr16 | 53,039,736 | G | A | 15 | YES | paternal | 0 | downstream | 0 |
| chr16 | 53,469,407 | A | G | 15 | NO | maternal | 1 | intergenic | 0 |
| chr16 | 67,910,704 | G | A | 15 | YES | paternal | 0 | intergenic | 0 |
| chr16 | 72,704,235 | G | A | 24 | — | paternal | 1 | intergenic | 1 |
| chr16 | 74,840,146 | G | A | 11 | — | maternal | 1 | intergenic | 1 |
| chr16 | 75,461,684 | T | A | 20 | — | paternal | 1 | intronic | 0 |
| chr16 | 76,140,720 | G | C | 21 | — | — | 0 | exonic | 0 |
| chr16 | 76,553,769 | T | A | 24 | — | paternal | 1 | intergenic | 0 |
| chr16 | 83,181,271 | T | G | 23 | — | paternal | 0 | intergenic | 0 |
| chr17 | 5,183,196 | T | A | 23 | — | paternal | 0 | intergenic | 0 |
| chr17 | 10,163,743 | G | T | 11 | — | paternal | 1 | intergenic | 0 |
| chr17 | 12,554,127 | G | A | 24 | — | — | 0 | intergenic | 0 |
| chr17 | 15,163,742 | G | A | 21 | — | — | 0 | intergenic | 0 |
| chr17 | 15,649,641 | G | A | 18 | — | paternal | 1 | intergenic | 0 |
| chr17 | 17,242,721 | T | C | 9 | YES | paternal | 0 | intronic | 0 |
| chr17 | 21,800,318 | C | G | 23 | — | — | 1 | intergenic | 0 |
| chr17 | 29,242,605 | C | T | 21 | — | — | 0 | intronic | 0 |
| chr17 | 34,910,084 | T | A | 19 | — | maternal | 0 | intergenic | 0 |
| chr17 | 36,964,951 | A | T | 11 | — | maternal | 0 | intergenic | 0 |
| chr17 | 45,470,220 | C | T | 23 | — | paternal | 1 | intergenic | 0 |
| chr17 | 48,484,121 | C | T | 8 | YES | paternal | 0 | intergenic | 1 |
| chr17 | 71,086,707 | G | A | 9 | YES | — | 0 | intergenic | 1 |
| chr18 | 74,909 | G | C | 14 | — | paternal | 0 | intergenic | 0 |
| chr18 | 7,623,725 | G | A | 14 | — | — | 0 | intergenic | 0 |
| chr18 | 10,368,083 | G | A | 23 | — | maternal | 0 | intronic | 0 |
| chr18 | 16,695,947 | T | G | 23 | — | paternal | 1 | intergenic | 0 |
| chr18 | 20,254,392 | G | T | 15 | NO | paternal | 1 | intergenic | 0 |
| chr18 | 25,356,508 | T | C | 21 | — | paternal | 1 | intergenic | 0 |
| chr18 | 26,897,367 | T | A | 22 | — | paternal | 0 | exonic | 0 |
| chr18 | 29,021,978 | G | T | 18 | — | — | 1 | intergenic | 0 |
| chr18 | 35,397,173 | T | G | 18 | — | maternal | 0 | intergenic | 0 |
| chr18 | 41,272,754 | G | A | 21 | — | — | 0 | intergenic | 1 |
| chr18 | 51,864,527 | T | G | 18 | — | maternal | 0 | intergenic | 0 |
| chr18 | 83,380,213 | A | G | 12 | YES | paternal | 0 | intergenic | 0 |
| chr18 | 88,184,224 | C | T | 22 | — | — | 0 | intergenic | 0 |
| chr18 | 91,802,898 | G | A | 22 | — | — | 0 | intergenic | 1 |
| chr19 | 15,405,188 | G | A | 20 | — | maternal | 0 | intergenic | 1 |
| chr19 | 17,579,029 | G | A | 23 | — | maternal | 0 | intergenic | 0 |
| chr19 | 20,884,800 | T | G | 23 | — | — | 0 | intergenic | 0 |
| chr19 | 28,102,224 | G | A | 21 | — | paternal | 0 | intergenic | 0 |
| chr19 | 28,795,346 | A | T | 19 | — | paternal | 1 | intergenic | 0 |

| | | | | | | | | | |
|---|---|---|---|---|---|---|---|---|---|
| chr19 | 35,345,935 | A | C | 11 | — | — | 1 | intergenic | 0 |
| chr19 | 40,911,797 | C | T | 24 | — | — | 1 | intergenic | 0 |
| chr19 | 41,809,167 | G | A | 25 | — | paternal | 1 | intergenic | 0 |
| chr19 | 47,453,854 | C | T | 22 | — | — | 1 | intergenic | 0 |
| chr19 | 52,628,816 | T | G | 23 | — | paternal | 1 | intergenic | 0 |
| chr19 | 55,616,865 | G | T | 14 | — | paternal | 1 | intergenic | 0 |
| chr19 | 57,763,280 | T | G | 23 | — | — | 0 | intronic | 0 |
| chr19 | 61,297,799 | T | G | 23 | — | paternal | 0 | intergenic | 0 |
| chr20 | 158,668 | T | C | 19 | — | paternal | 0 | intergenic | 0 |
| chr20 | 3,513,140 | A | T | 23 | — | paternal | 0 | intergenic | 0 |
| chr20 | 5,964,744 | A | G | 9 | NO | paternal | 0 | intronic | 0 |
| chr20 | 7,271,227 | T | G | 23 | — | — | 0 | intergenic | 0 |
| chr20 | 22,205,361 | C | T | 20 | — | paternal | 0 | intergenic | 1 |
| chr20 | 43,756,236 | C | T | 23 | — | paternal | 0 | intergenic | 0 |
| chr20 | 45,111,511 | A | G | 21 | — | maternal | 0 | intergenic | 0 |
| chr20 | 45,499,347 | A | C | 23 | — | — | 0 | intronic | 0 |
| chr20 | 52,420,953 | G | C | 23 | — | paternal | 0 | intronic | 0 |
| chr20 | 53,720,086 | T | G | 18 | — | paternal | 0 | intergenic | 0 |
| chr20 | 65,949,593 | T | C | 11 | — | — | 0 | intergenic | 0 |
| chr20 | 81,347,209 | G | A | 21 | — | paternal | 1 | intergenic | 0 |
| chr20 | 87,731,528 | G | A | 19 | — | — | 1 | downstream | 1 |
| chr20 | 88,434,883 | T | C | 25 | — | — | 0 | intergenic | 0 |
| chr20 | 88,622,035 | G | A | 15 | YES | — | 0 | intergenic | 0 |
| chr20 | 88,995,214 | C | T | 23 | — | — | 0 | intergenic | 0 |
| chr20 | 91,088,052 | G | C | 15 | YES | — | 1 | intergenic | 0 |
| chr21 | 4,270,552 | C | A | 11 | — | paternal | 0 | intergenic | 0 |
| chr21 | 5,871,346 | C | T | 8 | YES | paternal | 0 | intergenic | 1 |
| chr21 | 9,055,017 | A | G | 25 | — | maternal | 1 | intergenic | 0 |
| chr21 | 14,199,611 | T | C | 15 | NO | — | 0 | intergenic | 0 |
| chr21 | 17,254,412 | G | A | 15 | YES | — | 1 | intergenic | 1 |
| chr21 | 22,393,139 | C | G | 23 | — | — | 0 | intergenic | 0 |
| chr21 | 30,670,222 | G | A | 19 | — | paternal | 1 | intergenic | 0 |
| chr21 | 33,098,794 | T | G | 25 | — | — | 0 | intergenic | 0 |
| chr21 | 53,851,720 | A | C | 11 | — | maternal | 0 | intergenic | 0 |
| chr21 | 68,222,216 | G | A | 14 | — | — | 1 | intergenic | 1 |
| chr21 | 70,946,513 | A | C | 8 | NO | paternal | 0 | intergenic | 0 |
| chr22 | 14,126,689 | G | T | 8 | YES | maternal | 1 | intergenic | 0 |
| chr22 | 17,483,001 | C | T | 19 | — | paternal | 0 | intergenic | 1 |
| chr22 | 20,143,005 | G | T | 23 | — | maternal | 1 | intergenic | 0 |
| chr22 | 23,887,690 | G | A | 25 | — | — | 1 | intronic | 0 |
| chr22 | 25,482,344 | A | C | 9 | NO | — | 0 | intronic | 0 |
| chr22 | 30,262,434 | T | A | 15 | NO | — | 0 | intergenic | 0 |
| chr22 | 32,208,180 | G | A | 22 | — | — | 0 | intergenic | 1 |
| chr22 | 36,446,355 | T | C | 11 | — | paternal | 1 | intronic | 0 |
| chr22 | 38,703,441 | T | C | 25 | — | — | 0 | intergenic | 0 |

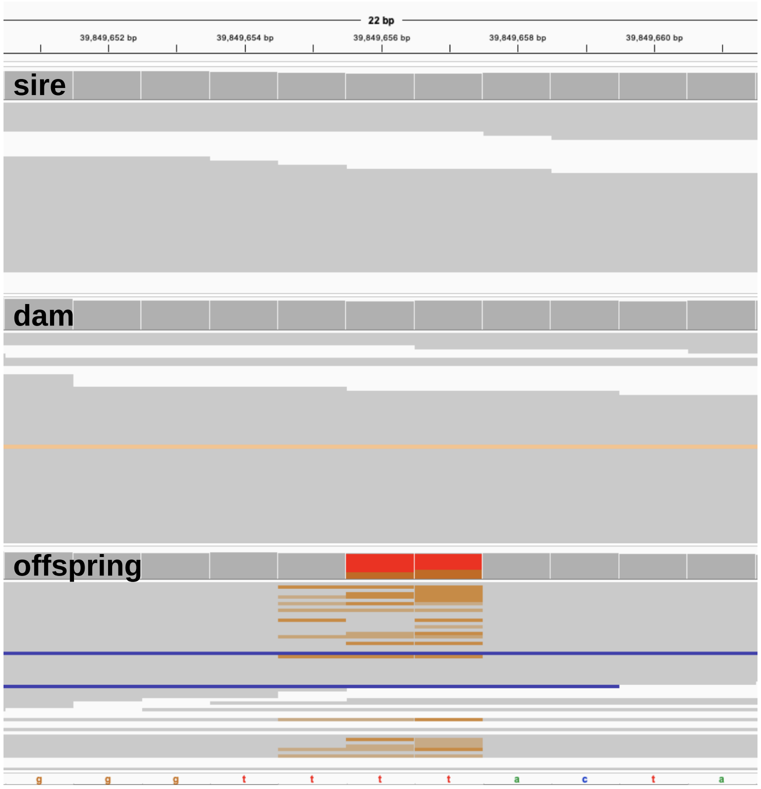

**Supplementary Figure 1**. Example of false positives associated with systematic genotyping errors observed in the vicinity of homopolymeric genomic tracts.

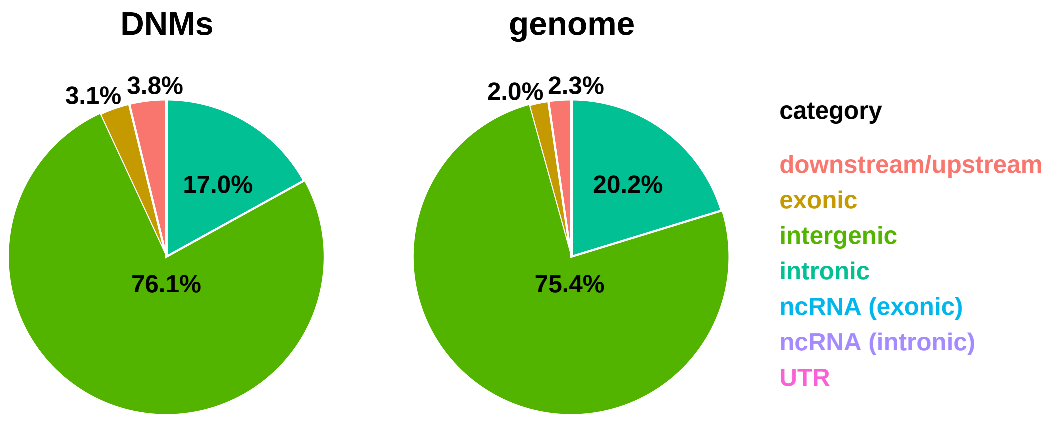

**Supplementary Figure 2**. Distribution of DNMs and genomic sites accessible to the study by genomic context based on the gene models available for the coppery titi monkey genome (GenBank accession number: GCA_040437455.1; Pfeifer et al. 2024).

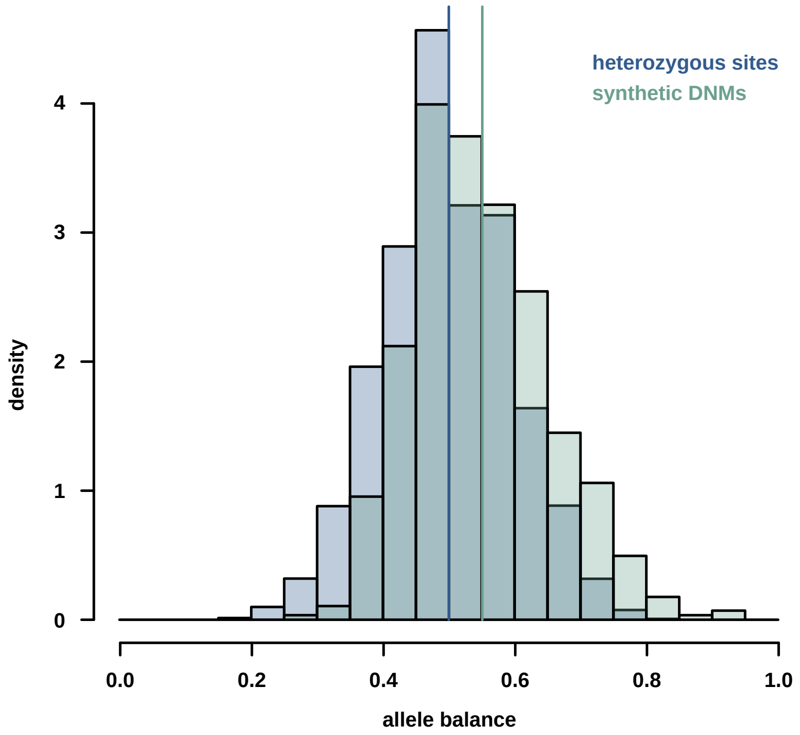

**Supplementary Figure 3**. Distribution of allele balance (i.e., the ratio of reads carrying the alternate vs reference alleles) observed for synthetic DNMs (shown in teal) and genuine heterozygous sites (blue).
